# Flexible control of speed of cortical dynamics

**DOI:** 10.1101/155390

**Authors:** Jing Wang, Devika Narain, Eghbal A. Hosseini, Mehrdad Jazayeri

## Abstract

Musicians can perform at different tempos, speakers can control the cadence of their speech, and children can flexibly vary their temporal expectations of events. To understand the neural basis of such flexible timing, we recorded from the medial frontal cortex of primates trained to produce different time intervals with different effectors. The activity of neurons was heterogeneous, nonlinear and complex. However, responses were unified under a remarkable form of invariance: firing rate profiles were temporally stretched for longer intervals and compressed for short ones. At the network level, this phenomenon was evident by flexible changes in the speed with which the population activity traced an invariant trajectory. To identify the origin of speed control, we recorded from both downstream caudate neurons and thalamic neurons projecting to the medial frontal cortex. Speed adjustments were a prominent feature in the caudate but not in the thalamus suggesting that this phenomenon originates within cortical networks. To understand the underlying mechanisms, we created recurrent neural network models at different levels of complexity that could explain flexible timing with speed control. Analysis of the models revealed that the key to flexible speed control was the action of an external input upon the nonlinearities of individual neurons whose recurrent interactions set the network’s relaxation dynamics. These findings demonstrate a simple and general mechanism for conferring temporal flexibility upon sensorimotor and cognitive functions.

---

Mental capacities such as anticipation, motor coordination, deliberation, and imagination lie at the core of higher brain function. A fundamental feature of these capacities is that they are not tied to immediate sensory or motor events and unfold at different timescales. To support such temporal flexibility, the brain must control the dynamics of ongoing patterns of neural activity. However, the mechanisms that control the dynamics are not known. A simple behavior where flexible adjustments of dynamics are required is motor timing; i.e., controlling when to initiate a movement. To test the neural basis of temporal flexibility in motor timing, we developed a task in which monkeys used different contextual cues to produce either a 800 ms or a 1500 ms interval, responding either by saccade or manual button press (Fig. 1a).

**Fig. 1.**
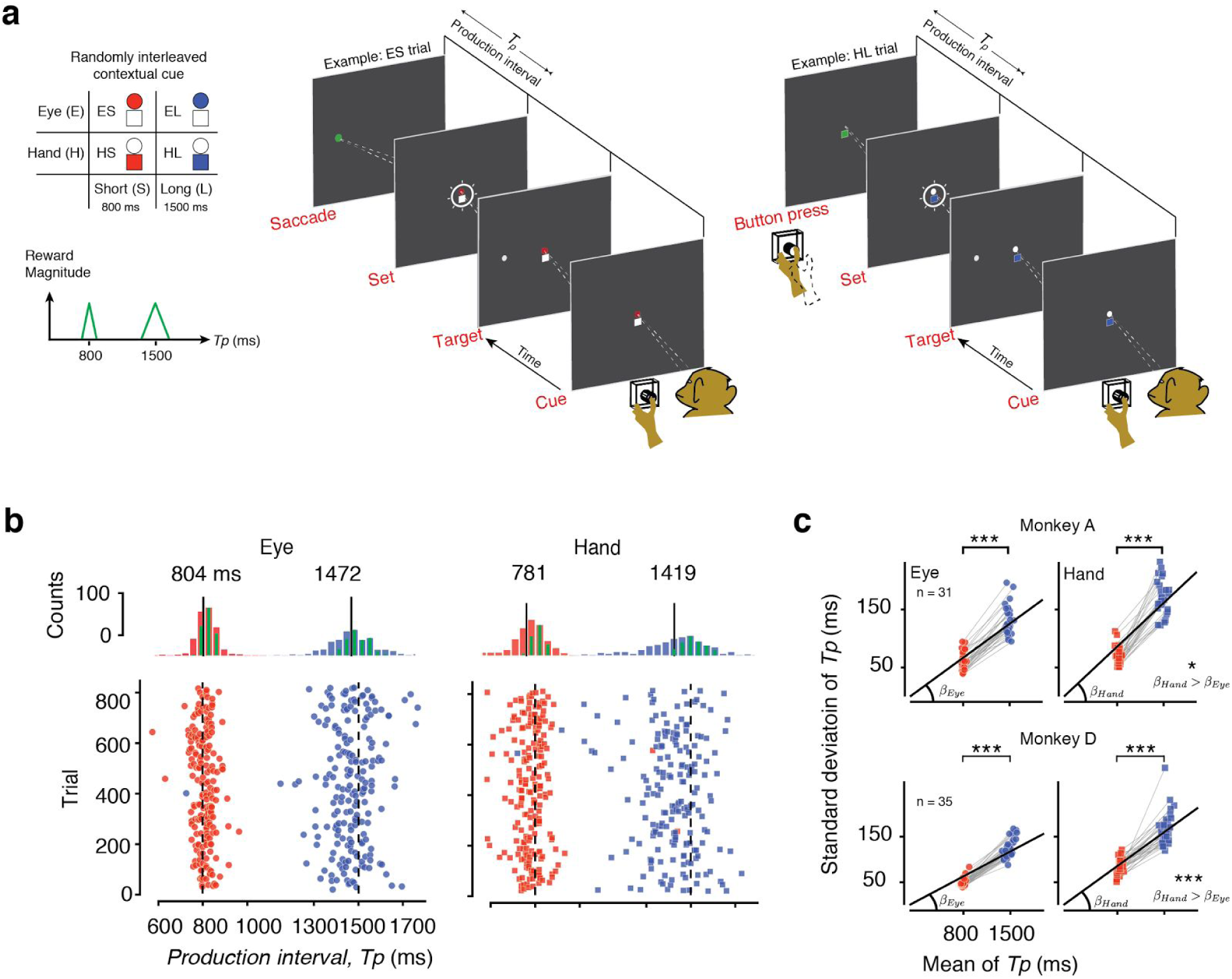
Time production task and behavior. **(a)** Trial structure. Animals had to produce either a 1500 ms (*Long*), or an 800 ms (*Short*) interval, either by making a saccade (*Eye*) or a button press (*Hand*). These four conditions were randomly interleaved and were cued throughout the trial by the color and shape of two central stimuli, a circular fixation for the eye and a square that cued the animal to place its hand on a button. The colored shape (circle or square) cued the effector, and the hue (red or blue) cued the desired interval (red for *Short* and blue for *Long*). After a random delay (0.5 - 1.5 sec, uniform hazard), a filled white circle was flashed to the left or right of the fixation point. This peripheral flash specified the saccadic target for the eye trials and played no role in the hand trials. After another random delay (0.5 -1.5 sec, uniform hazard), a *Set* cue (a ring flashed around the two fixation stimuli) initiated the motor timing epoch. The animal’s production interval (*Tp*) was measured as the interval between Set and when either the saccade was made or the button was pressed. The saccade was rewarded if it was directed to the memorized location of the peripheral target. When *Tp* for the desired effector (eye or hand) was within a specified reward window, the peripheral target (or the square fixation) turned green, auditory feedback was provided, and animal was rewarded with juice. The reward window was set adaptively on a trial-by-trial basis and independently for the *Short* and *Long* conditions so that the animal received reward on approximately 50% of trials both both interval context on every session (on average, 57% in monkey A and 51% in monkey D). The reward magnitude increased linearly with accuracy as shown by the green triangular reward function. Two example trials, one for the *Eye/Short* (ES) condition (left) and one for the *Hand/Long* (HL) condition are shown. **(b)** A typical behavioral session showing *Tp* while the animal flexibly switched between the four trial conditions. For clarity the *Eye* (left) and *Hand* (right) trials are plotted separately although during the task they were randomly interleaved. The top 4 histograms show the distribution of *Tp* for each condition with rewarded trials in green. The vertical lines correspond to the mean values that are also reported numerically. **(c)** Top: Behavior across 31 sessions for animal A. The standard deviation of *Tp* scaled with its mean for both *Eye* (left panel, circle) and *Hand* (right panel, square) and both intervals (red and blue). The mean ± s.e.m of *Tp*s was ES: 810 ± 48.9 ms, EL: 1495 ± 117 ms, HS: 822.3 ± 53.7 ms, HL: 1486 ± 136 ms. The variability was significantly higher for the *Long* compared to the *Short.* The average Weber fraction (ratio of standard deviation to mean) for the *Hand* (*β_Hand_*) was significantly larger that *Eye* (*β_Eye_*) (one-tailed paired sample *t*-test, n = 31). Bottom: Same as the top panel for animal D. The mean ± s.e.m of *Tp*s for the 4 trial conditions were: ES: 808 ± 56.1 ms, EL: 1481 ± 137 ms, HS: 836.7 ± 91.3 ms HL: 1521 ± 177 ms. The variability was significantly higher for the *Long* compared to the *Short* and, *β_Hand_* was significantly larger than *β_Eye_* (one-tailed paired sample *t-*test, n = 35) *α* = .05 (*) and *α* = .001 (***) levels.

While monkeys performed the task, we recorded neural activity in the dorsomedial frontal cortex (MFC), which has been implicated in the inhibition ^1,2^, initiation ^3–5^, and coordination ^6–12^ of movements. Consistent with previous work in humans ^13–19^, nonhuman primates (NHPs)^3,20–26^ and rodents ^27–32^, MFC responses were modulated by elapsed time. The activity of a large proportion of neurons exhibited an intriguing form of temporal invariance: responses were stretched or compressed in accordance with the produced interval. Temporal scaling at the level of single neurons is equivalent to an adjustment of speed with which the population activity evolves along an invariant trajectory. Two properties made this change of speed noteworthy. First, unlike previous work where temporal scaling was reported during adaptation and learning ^33,34^, here, speed control constituted rapid trial-by-trial switches based on contextual cues, which cannot be explained by slow plasticity mechanisms. Second, the majority of neurons that exhibited temporal scaling had nonlinear and heterogeneous response profiles making the scaling phenomenon inconsistent with existing models of timing ^35,36^ including pacemaker accumulator ^37–40^, oscillations ^41–43^ liquid-state machine population clocks ^44–48^. These considerations necessitated a new perspective that could explain temporal scaling in neurons with highly complex response profiles.

As a first step toward investigating the relevant neural substrates, we examined temporal scaling in the caudate nucleus downstream of MFC and regions of the thalamus projecting to MFC that have been implicated in motor timing ^20,22,23,25,49–52^. Results were best explained by the hypothesis that speed control originated in the cortex.

To investigate the potential underlying mechanisms, we analyzed the dynamics of recurrent neural network models that were either engineered or trained to produce different time intervals using context-dependent inputs. By setting the models in certain input-output regimes, we discovered a simple and novel mechanism that enabled the rapid and flexible adjustment of the speed of dynamics in networks of heterogeneous neurons.

## Behavior

On each trial, the color and shape of the fixation point served as a contextual cue (‘*Cue*’) indicating the desired interval and response effector respectively (Fig. 1a). We refer to the trial conditions as EL, ES, HL and HS, where E and H indicate *Eye* and *Hand*, and S and L indicate *Short* (800 ms) and *Long* (1500 ms) intervals. Production intervals (*Tp*) were measured from the time of a brief ‘*Set*’ flash until the time of movement initiation (‘*Saccade*’ or ‘*Button press*’). The four conditions were randomly interleaved and animals were able to successfully switch between conditions on a trial-by-trial basis (Fig. 1b). They produced accurate *Tp*s whose variability increased for the *Long* condition compared to *Short* (Fig. 1c). This is consistent with the Weber’s law and is a well-known property of timing behavior ^53,54^. The Weber fraction of *Tp*s across the two contexts (ratio of standard deviation to mean) was significantly larger for button presses compared to saccades (one-tailed paired sample *t*-test, for monkey A, n = 31 sessions, *P* < .05, and for monkey D, n = 35 sessions, *P* < .001).

## Causal experiments and single-unit electrophysiology

We first verified that the regions of interest in MFC (Fig. 2a) were causally involved in this task (Fig. 2b). Reversible inactivation with muscimol (GABA_A_ agonist) significantly impaired performance for both *Long* and *Short* intervals as measured by the distribution of within-session increases in root-mean-squared error (RMSE) after the muscimol injection, when compared to before (Table 1). The drop in performance in each session comprised of changes in both mean and standard deviation (Fig. 2b). However, no significant impairment was measured after saline injection (Table 1). Furthermore, muscimol inactivation had no significant effect on reaction times during a memory saccade task (Table 1). Based on these results, we concluded that MFC played a causal role in the main motor timing task ^3,30,32^.

**Fig. 2.**
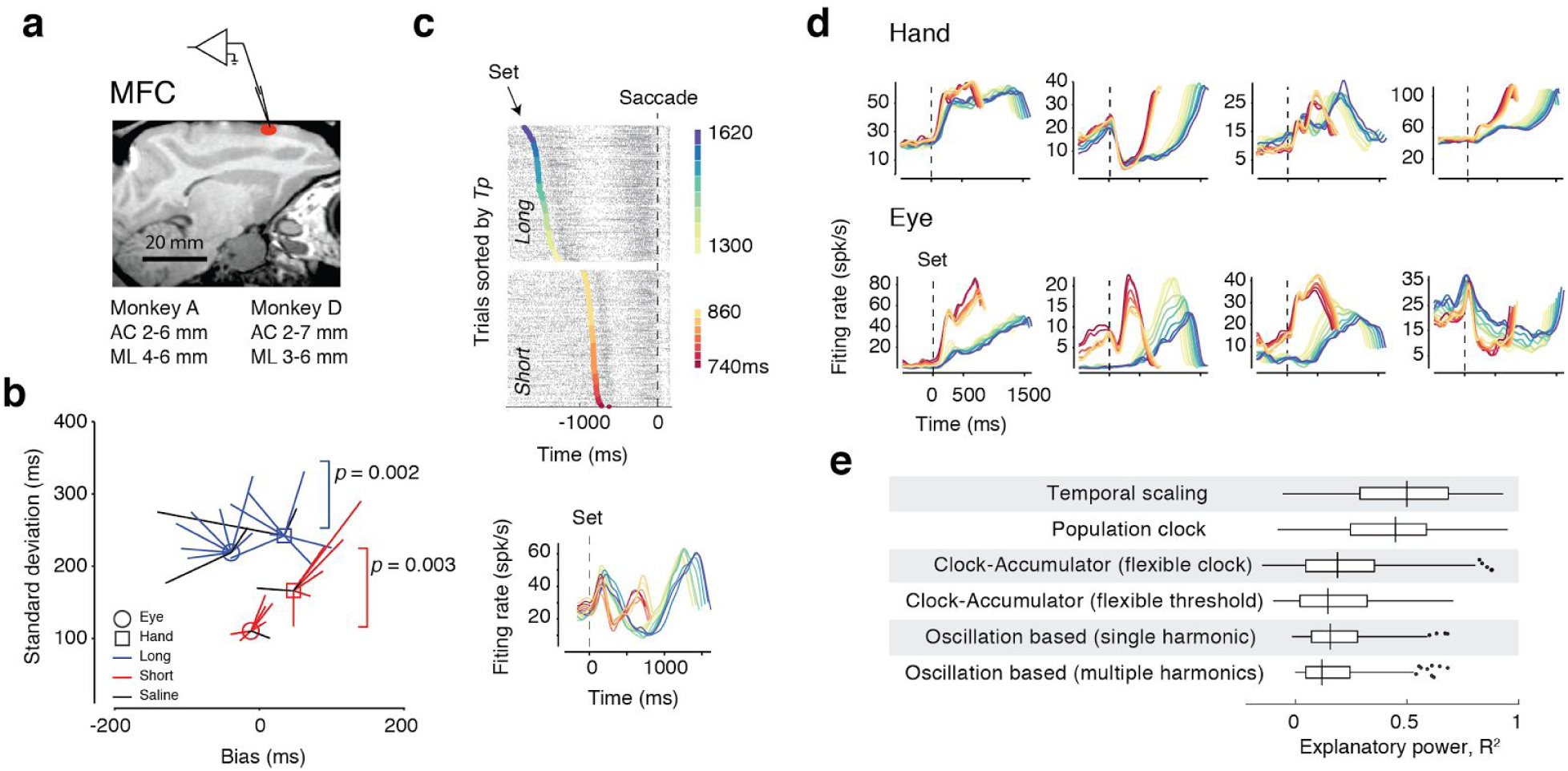
Medial frontal cortex inactivation and electrophysiology. **(a)** Parasagittal view of one of the animals (monkey D) with a red ellipse showing the region targeted for inactivation and electrophysiology. Bottom: the corresponding stereotactic coordinates in each animal with respect to anterior commissure (AC) and midline (ML). **(b)** Muscimol inactivation. Each line in each panel shows the change in bias (abscissa) and change in standard deviation (ordinate) after muscimol injection in one behavioral session separated for *Hand* (square) and *Eye* (circle). Red and blue correspond to *Short* and *Long* trials, and black, to Saline control sessions. The reported p-values correspond to a test of whether inactivation increased RMSE across sessions (RMSE^2^ = Σ(*Tp* - *Ts*)^2^ = Bias^2^ + Var, one-tailed paired t-test, see Table 1). No significant change of RMSE was observed after saline injection (Table 1). **(c)** Computing post-stimulus-time histograms (PSTHs). Top: Raster plot of spike times (black ticks) for a single neuron aligned to movement initiation time (dashed line) across trials (rows) for an example neuron. Trials were sorted and grouped into bins (colors) according to the produced interval (*Tp*). Bottom: PSTHs that were computed with respect to movement initiation time were subsequently plotted aligned to the time of Set (dashed line). The Set across trials in the top panel, and the activity profiles in the bottom panels were colored according to *Tp* bins (legend). **(d)** Activity profile of 8 example neurons for *Hand* (top) and *Eye* (bottom) conditions computed as described in (c). **(e)** Analysis of single neurons with respect to various model of timing. Whisker plot (median: center line; box: 25th to 75th percentiles; whiskers: ± 1.5 × the interquartile range; dots: outliers) showing the range of R^2^ values captured by six models fitted to the PSTHs of individual neurons. The “Temporal scaling” model (top) had the highest explanatory power of R^2^ in comparison with all other models (one-way ANOVA, F_6, 2444_ = 156.5, *P* < 0.001, and one-tailed paired sample *t*-test between Temporal scaling’ and ‘Population clock’ model, n = 416, *P<* .001).

**Table 1.** Number of neurons recorded in each area in each animal

|  | MFC<br>(included/total) | Caudate<br>(included/total) | Thalamus<br>(included/total) |
| --- | --- | --- | --- |
| Monkey A | 281/356 | 101/200 | 481/534 |
| Monkey D | 135/166 | 177/309 | 365/402 |
| Both animals | 416/522 | 287/509 | 846/936 |

## Temporal scaling of complex response profiles

To characterize the post-stimulus-time-histogram (PSTH) of MFC neurons in anticipation of the motor response, we binned trials based on production intervals and computed average firing rates for each bin from spike counts after aligning trials to the time of the motor response (Fig. 2c). Neurons had complex and heterogeneous response dynamics including linear, nonlinear, monotonic, non-monotonic and even multi-peaked activity profiles (Fig. 2d). This diversity could not be accounted for by existing models of timing. To quantify this, we tested single-neuron PSTHs against predictions of various models of motor timing using a cross-validation procedure (Fig. 2e). We considered two variants of the clock-accumulator model, one in which flexible timing was achieved by adjusting a threshold over a ramping process, and one in which the clock was adjusted. Since clock models can only accommodate neurons with linear ramping profiles ^37–40,55^ it was not surprising that both clock-accumulator models were unable to explain the nonlinear profiles exhibited by much of the population. This result was substantiated by fitting polynomials of different degrees to PSTHs using a cross-validation procedure (see Methods). This exercise revealed that only 11% (47/416) of neurons could be explained by the clock model and the remaining neurons had complex and nonlinear response profiles that deviated significantly from linear ramping. This number was robust to changes in the formulation of the clock-model. For example, making the clock model more flexible by allowing the starting and terminating points of the ramp to vary by up to 200 ms increased the number of neurons explained by this model by only 4%. We also tested two oscillation-based models of interval timing, in which the response time is determined by the collective phase of multiple oscillators with different frequencies ^41–43^. In one variant, a single sinusoid was fit to the response of each neuron, and in another, multiple sinusoids (up to 4) of different frequencies were used. These models were also unable to capture the diversity of MFC responses (Fig. 2e). Finally, we tested MFC responses against a relatively unconstrained population-clock model ^44–48^. In this model, each neuron is allowed to have a unique PSTH, and the movement is triggered when a decoder detects an interval-dependent pattern of activity across the neurons. Accordingly, we modeled each neuron by the best-fitting polynomial that could capture the activity profile in both the *Short* and *Long* contexts. This model performed better than both the clock-accumulator and oscillation models owing to the higher degrees of freedom associated with polynomial fits. However, MFC data did not fit with a key qualitative prediction of the population clock model. In this model, neurons are expected to have identical PSTHs for the *Short* and *Long* conditions for the entire duration of the timing interval. However the vast majority of neurons in MFC had PSTHs that were distinct for the *Short* and *Long* conditions(Fig. 2d).

Our initial inspection of the difference between response dynamics in *Short* and *Long* conditions suggested that PSTHs for different bins had a high degree of self similarity when stretched or compressed in accordance with the produced interval (Fig. 2c,d). This was true both for fluctuations of produced intervals within each temporal context (i.e., 800 ms or 1500 ms), and across the two temporal contexts. To verify this observation quantitatively, we asked whether the PSTHs were more accurately captured by a temporally-scaled polynomial function. This formulation clearly outperformed all other models in terms of explanatory power (Fig. 2e, one-way ANOVA, F_6, 2444_ = 156.5, *P <* 0.001). Note that the clock-accumulator with a flexible clock also exhibits temporal scaling but it cannot capture scaling in neurons with complex response profiles, which as we reported, comprised 89% of task-modulated MFC neurons. These observations highlighted the need for an alternative model that could explain temporal scaling across neurons with such heterogeneous response profiles.

## Speed control across the population

The phenomenon of temporal scaling at the level of single neurons has an intuitive interpretation at the population level. When the population activity is plotted within a coordinate system in which each axis corresponds to the firing rate of a single neuron, also known as the *state space* ^56^, response dynamics can be depicted as a point (i.e., neural state) moving along a trajectory. In this representation, temporal scaling corresponds to changing the speed with which the neural state traverses a single trajectory in high-dimensional space.

If all the responses were scaled perfectly, neural trajectories associated with different intervals would be identical. However, for many neurons, firing rate modulations did not scale perfectly (Fig. 2d). We quantified the degree of scaling by a *scaling index* (SI) that was computed as a coefficient of determination (R^2^) across temporally-scaled PSTHs associated with different production interval bins. With this measure, we verified that neurons exhibited variable degrees of temporal scaling (Supplementary Fig. 1). The imperfect scaling was also evident by visualizing neural trajectories within the space spanned by the first three principal components (PCs). Within this space, neural trajectories did not overlap (Fig. 3a) suggesting that neural responses had features that did not scale perfectly across intervals.

**Fig. 3.**
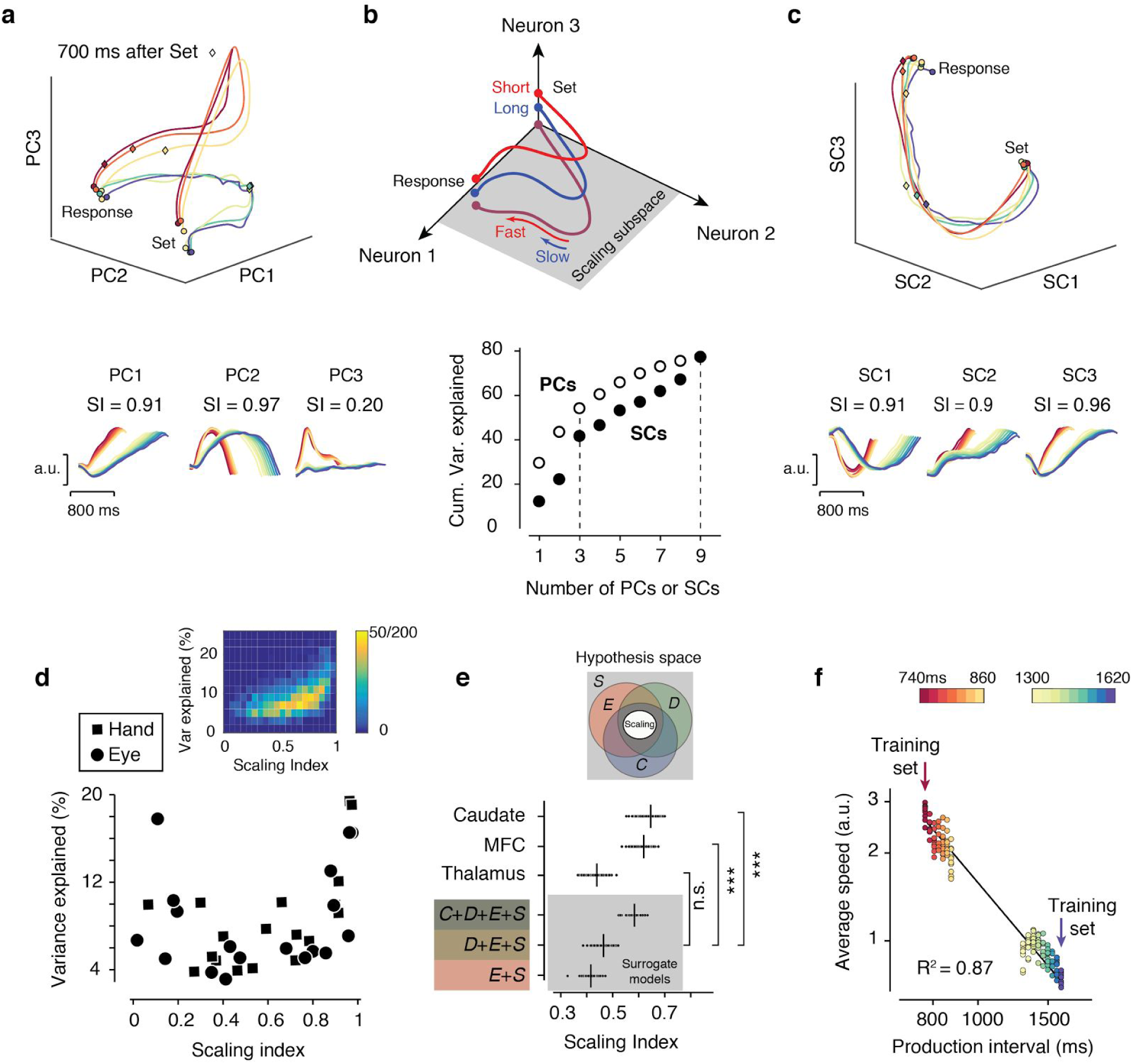
Temporal scaling in the medial frontal cortex at the population level. **(a)** Top: Population activity profiles for hand trials of one animal projected onto the first 3 principal components (PCs) from the time of Set to the time of button press (Response). Activity profiles associated with different produced intervals for *Short* and *Long* conditions are plotted in different colors (see color bar in Fig. 3f). The state at 700 ms after Set is shown along the trajectories (diamond). Bottom: The time course of the first three PCs with the corresponding scaling index (SI) values. **(b)** Top: Schematic drawing illustrating the scaling subspace. The response dynamics associated with a *Short* (red) and *Long* (blue) produced interval (*Tp*) are depicted as distinct trajectories in the state space. The start and end of the trajectory are demarcated by Set and Response (filled circles). Projections of neural responses onto a scaling subspace result in overlapping trajectories (purple) whose speed determines the produced interval (fast for short and slow for long). Bottom: Cumulative percentage variance explained by PCs and SCs. (c) Top: Population activity sorted according to *Tp* bins and projected onto the first 3 scaling components (SCs). As expected, in this subspace, the trajectories overlap. Bottom: The first three SCs with the corresponding SI values. Because of cross-validation, the scaling index of SCs of the test data were not in decreasing order, although they were for the training dataset (not shown). **(d)** Variance explained for individual SCs as a function of scaling index. SCs with the larger scaling indices explain a large percentage of variance for both *Hand* (square) and *Eye* (circle) conditions. Inset: An unbiased estimate of variance explained as a function of scaling index derived from random linear projections of the MFC activity in the state space. The plot includes 200 random projections binned and pseudocolored to indicate the frequency of occurrence. The data shows that high scaling indices are associated with high variance explained. **(e)** Comparison of scaling index between the MFC, caudate and thalamus with three surrogate models that capture the observed firing rate statistics with increasing levels of sophistication. The data for surrogate models were generated from Gaussian processes that were constrained to match the smoothness in the data (i.e., same smoothness, S). In the first model (*E+S*), surrogate data were additionally constrained such that responses for different production intervals were the same at the time of Set and the time of response (i.e., same endpoints, *E*). The second model (*D+E+S*) additionally constrained the surrogate data to have the same dimensionality as the MFC data (i.e., same dimension, *D*). The third model (*C+D+E+S*) further constrained the data such that the R^2^ correlation between responses to different produced intervals were the same as what would be expected from perfectly scaling responses (i.e., same correlation, *C*). Each model consisted of the same number of neurons as that in the MFC data, and the number of bootstrapped samples for each model was n = 200. The plot shows the the average scaling index across all SCs computed from bootstraps (small circles) along with the corresponding mean (vertical line) for each of three brain areas and each of the surrogate models. The average scaling index for each surrogate model was significantly lower than the values associated with the MFC and caudate, but not for the thalamus (one-tailed two-sample *t-*tests, n = 200, *** indicates *P* < .001, see main text). The inset shows the hypothesis space in relation to various constraints and their combinations with distinct colors and their overlaps. Perfect scaling (middle ellipse) is a subset of the possibilities that satisfy all four constraints. The colors correspond to those used to label surrogate models on the ordinate. **(f)** The speed of neural trajectory within the scaling subspace spanned by the first 3 SCs predicted average *Tp*s across bins.

We hypothesized that perfect speed control might emerge within a subspace where the projection of neural trajectories are invariant; i.e., *scaling subspace* (Fig. 3b). As a first step, we examined the degree of scaling in a subset of PCs. Using the same SI metric used for single neurons, we found that the first two PCs that explained nearly 40% of the variance (Fig. 3b, bottom) had a scaling index of 0.91 and 0.97, respectively (Fig. 3a, bottom). The third PC, however, did not exhibit temporal scaling and had a scaling index of 0.20. This provided evidence that speed control was present within a subspace whose dimensions explained a large percentage of variance (i.e., PC1 and PC2).

However, the dimensions of scaling do not have to perfectly coincide with PCs. To specifically target the dimensions of scaling, we developed a novel procedure to find a subspace in which trajectories were invariant and the activity was governed by speed control. Analogous to PCs that are ordered with respect to variance explained, we sought components that were ordered according to the degree of temporal scaling in the data. We formulated this as a cross-validated optimization problem that searched for dimensions in which the distance between neural trajectories of different temporal durations was minimized (see Methods). This method furnished a set of *scaling components* (SCs) that were ordered according to the degree of scaling in the data (Fig. 3c). Compared to PCs, SCs explained less variance suggesting that the scaling dimensions were not identical to PC dimensions, which capture maximum variance by design.

The SI values were relatively large for the first few SCs indicating that the optimization process correctly identified the scaling dimensions. (Fig. 3c, bottom, Supplementary Fig. 2). When responses were projected onto the subspace spanned by the first three SCs, they traced a nearly identical trajectory with different speeds (Fig. 3c, top), which is precisely what the scaling subspace hypothesis predicts. Note that because of cross-validation, the scaling index for SCs of the test data does not have to be in decreasing order (Fig 3c., bottom), although it was the case for the dataset which was used to determine the SCs (not shown).

Next, we asked how much of the variance of neural responses can the scaling subspace account for. To address this question, we performed two complementary analyses. First, we examined the relationship between the degree of scaling (SI) and the variance explained for each SC. SCs with large SIs explained a relatively large percentage of variance (Fig. 3d) suggesting that scaling was a prominent feature of neural activity during flexible time interval production. Second, we developed a procedure for quantifying the relationship between scaling and variance without relying on projections onto specific directions, such as PCs or SCs. We used a bootstrap procedure and quantified the relationship between variance explained and SI along 200 random projections in the state space. We then constructed a two-dimensional probability distribution of the relationship between variance explained and scaling across those random projections (Fig. 2d, inset). This analysis verified that the dimensions with large degrees of scaling also explained a large portion of the variance.

To further validate our conclusions, we examined the sufficiency of SI as a measure of scaling asking whether SI can be used as a reliable measure of temporal scaling. We tackled this problem by contraposition, asking whether the lack of scaling would result in small scaling indices. We used Gaussian processes to create non-scaling surrogate data that matched MFC responses in terms of smoothness, starting/terminal firing rates, dimensionality, and the correlation between *Short* and *Long* PSTHs (see Methods, and Supplementary Fig. 3). The surrogate data, despite being matched to the statistics of MFC responses, had relatively smaller SIs than computed for MFC neurons (Fig. 3e). This comparison to the non-scaling surrogate data verifies that a significant portion of variance in MFC resides within a scaling subspace in which activity evolves along invariant trajectories at different speeds.

Finally, we quantified the relationship between speed in the scaling subspace and behavior. We used a cross-validation procedure in which we derived the scaling subspace from a subset of shortest and longest trials, and asked whether the speed of neural trajectories of the remaining trials in that subspace could predict production intervals (*Tp*). This analysis provided strong evidence for the correspondence between the speed in the scaling subspace and *Tp*: longer *Tp*s were associated with slower speeds (Fig. 3f and Supplementary Fig. 4), and the average speed was inversely proportional to *Tp* (R^2^ = 0.87). These results establish speed control in the MFC scaling subspace as a key principle that governs behavioral variability within each temporal context, and enables the flexible control of motor timing across the two timing contexts.

## Speed control across cortico-basal ganglia circuits

Having established speed control in MFC as a potential mechanism for temporal flexibility, we asked whether this property was unique to MFC or whether it was also present in other upstream and downstream areas. We focused on various nodes of the cortico-basal ganglia circuits that have been implicated in motor timing ^20,22,23,25,49–52^. First, we investigated putative targets of MFC in the caudate ^57–59^. We found that the region of interest in the caudate (Fig. 4a) was causally involved in the motor timing task as evidenced by reversible inactivation (Table 1).

**Fig. 4.**
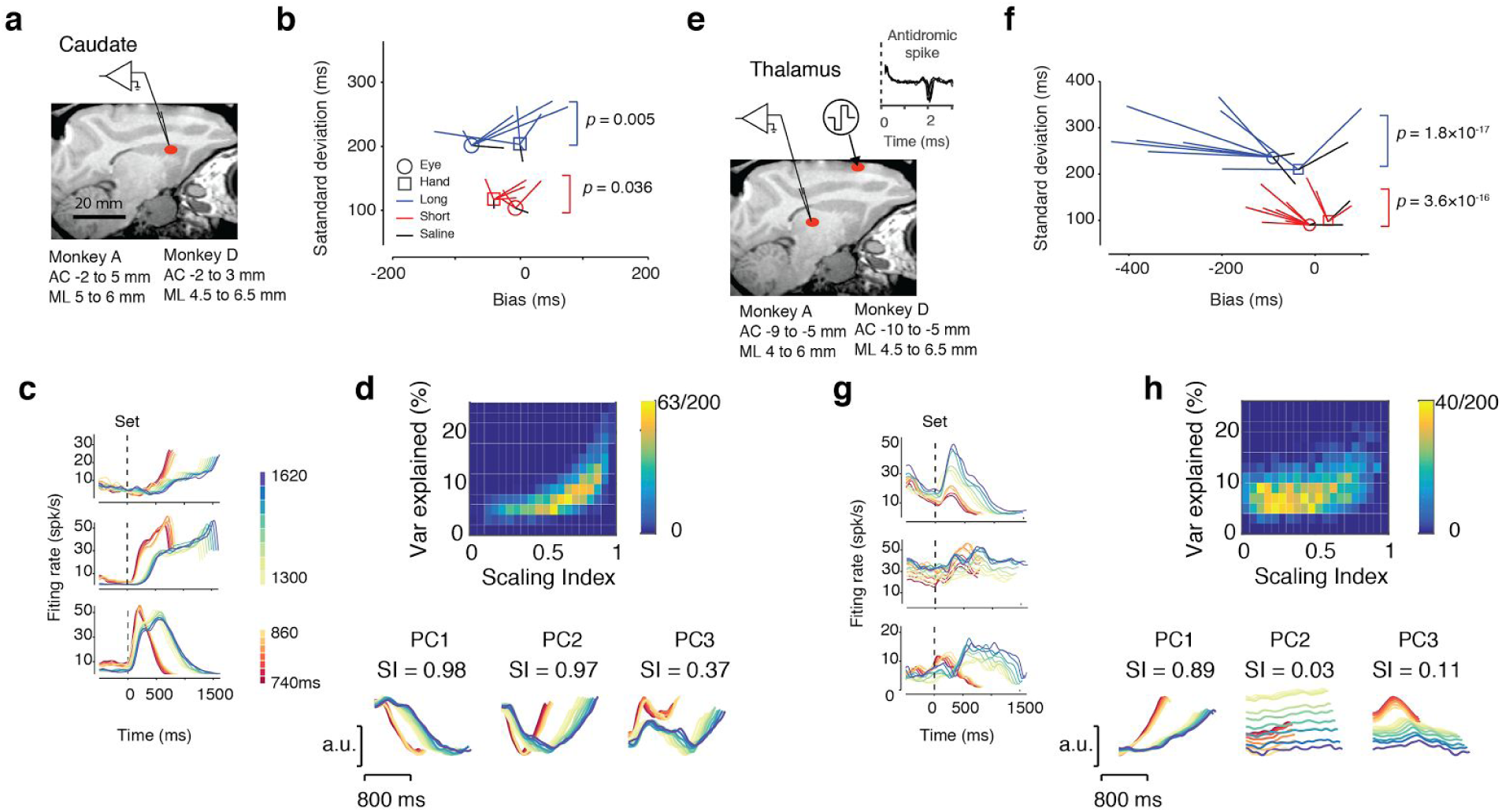
Inactivation, electrophysiology and temporal scaling in the caudate and thalamus. **(a)** Same as Fig. 2a with a red ellipse showing the region of caudate targeted for inactivation and electrophysiology. Bottom: stereotactic coordinates in each animal. **(b)** Muscimol inactivation in the caudate. Results are presented in the same format as in Fig. 2b. Statistics are reported in Table 1. **(c)** Activity profile of three example neurons across production time (*Tp*) bins (colors) computed as described in Fig. 2c. **(d)** Top: The relationship between variance explained and scaling index in the caudate. Results are presented in the same format as the inset of Fig. 3d. Bottom: The first three PCs with the corresponding scaling index (SI) values. **(e)** Same as panel a showing the region of interest in the thalamus. We recorded from neurons in the region where MFC-projecting neurons were identified antidromically. Inset: example of reliable and low-latency spikes detected after antidromic stimulation. **(f)** Muscimol inactivation in the thalamus. Results are presented in the same format as in Fig. 2b. Statistics are reported in Table 1. **(g)** Same as panel c for three example neurons in the thalamus. The example neuron in top right was antidromically identified. **(h)** Top: Variance explained as a function of SI in the thalamus (same format as in panel d). Results in the thalamus are qualitatively different from the caudate (panel d) and MFC (Fig. 3d) in that most projections in the state space do not exhibit temporal scaling. Bottom: The first three PCs and the corresponding SI values in the thalamus.

Electrophysiological recordings (Fig. 4c) indicated that, caudate responses, like those in MFC, were complex and heterogeneous, and PSTHs were different between *Short* and *Long* trials. We analyzed caudate responses in terms of temporal scaling and speed control. At the level of single neurons, the distribution of SIs was similar to MFC (Supplementary Fig. 1) suggesting that MFC neurons and their putative targets in the caudate exhibit similar degrees of scaling. At the population level, analysis of PCs and SCs verified the presence of a scaling subspace in the caudate (Fig. 3e, Supplementary Fig. 5). Finally, the SI values of PCs as well as an unbiased analysis of responses across random projections in the state space indicated that dimensions with strong scaling explained a large part of variance in the data (Fig. 4d). These analyses substantiated that neural signals in the caudate share the same key properties with MFC and may be part of the circuit involved in subspace speed control.

In addition to receiving inputs from MFC, the basal ganglia also projects back to MFC through the thalamus. The presence of this anatomical substrate raises the possibility that MFC inherits temporal scaling from the basal ganglia via transthalamic projections. To test this possibility, we examined neural activity in a region of the thalamus where MFC-projecting thalamocortical neurons were identified antidromically (Fig. 4e; see Methods). Consistent with previous work ^52^, reversible inactivation indicated that this area played a causal role in timing behavior (Table 1). However, several observations indicated that the function of thalamocortical signals was different from that of the caudate and MFC (Fig. 4g). First, SIs of single thalamic neurons were significantly smaller across the population compared to the other areas (n_MFC_ = 416, n_Cd_ = 278, n_thalamus_ = 846, one-tailed two sample *t-*tests, *P* < .001, see Supplementary Fig. 1). Second, scaling in the thalamus was significantly smaller than the C+D+E+S surrogate data (one-tailed two sample *t*-test, n = 200, *P* < .001 comparing to C+D+E+S surrogate model, Fig. 3e). Third, scaling was less prominent in the thalamus as indicated by the relationship between the magnitude of scaling and variance explained along random projections in the state space (Fig. 4h). Fourth, unlike the caudate and MFC, neural trajectories in the thalamus were not invariant in the space spanned by the first three SCs (Supplementary Fig. 5). This was also evident in the profile of the second PC whose relationship to different *Tp*s was a systematic shift in average value – i.e., not scaling. Together, these observations provide strong evidence that thalamic neurons exhibit significantly less scaling than the MFC neurons they project to. Since the output of the basal ganglia to cortex is routed through the thalamus, the weak scaling in thalamocortical neurons implicates that the scaling subspace is likely to originate within MFC or other cortical circuits projecting to MFC.

## A model for flexible subspace speed control

Since the timescales of MFC response modulations were slower than the intrinsic time constants of single neurons, we assumed that the observed dynamics were the result of network-level interactions. Motivated by recent advances in understanding the dynamics of motor, premotor and prefrontal cortical areas using recurrent network models ^60–62^, we trained a recurrent neural network model with random connectivity to flexibly produce different time intervals in response to context inputs (Cue) whose magnitude specified the desired interval (Fig. 5a). To mimic the task of the monkey, at a random time after the Cue onset, the network was administered a transient pulse (Set), which signaled the start of the time interval. The network was trained to generate a single output (a weighted linear sum of its units) that would initiate a “response” when the output breached a fixed threshold ^63,64^.

**Fig. 5.**
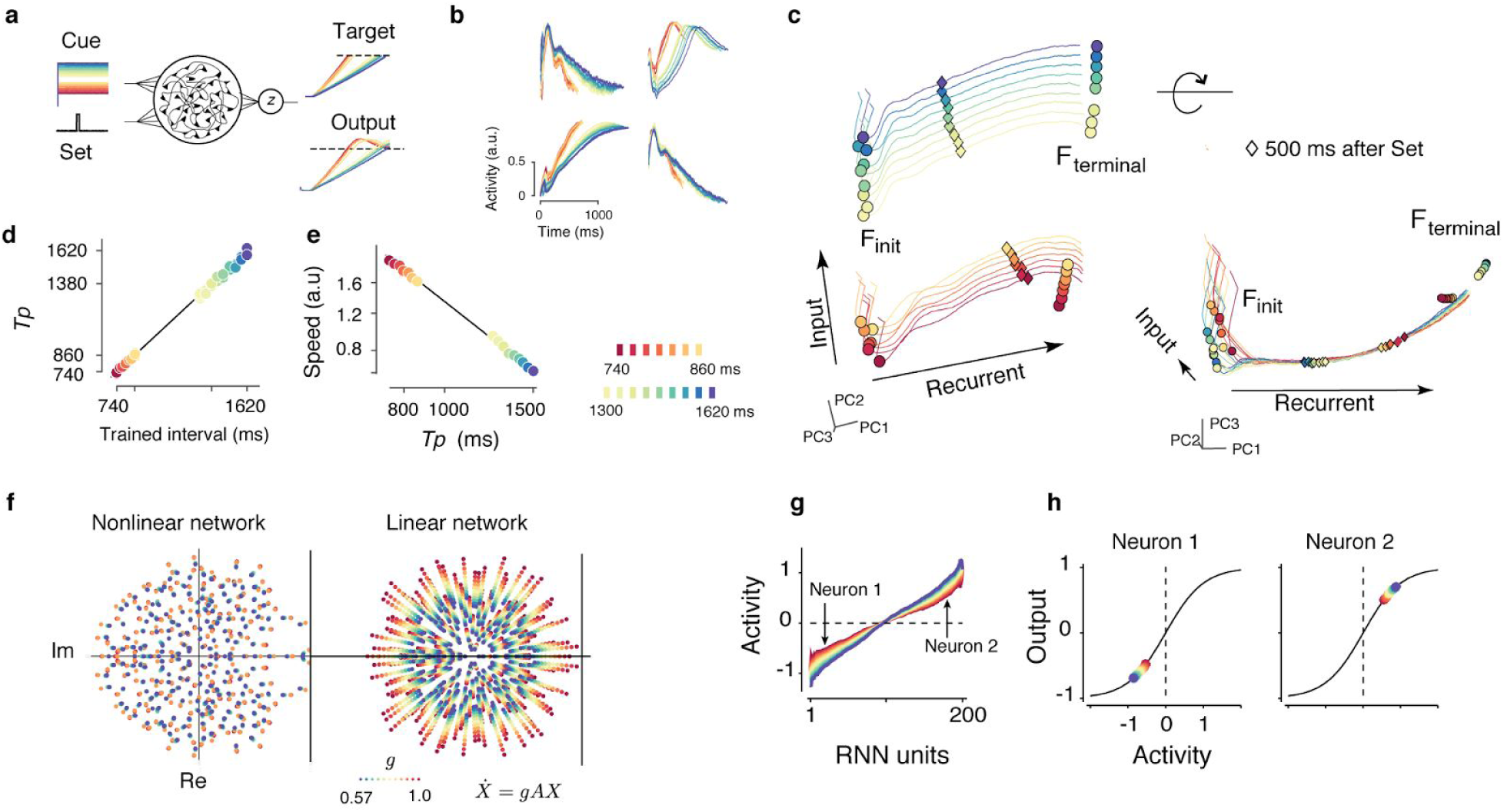
Recurrent neural network model dynamics. **(a)** A recurrent neural network model that receives an input (Cue) whose strength depends on the desired interval (different colors), and a transient Set pulse that initiates the timing interval. After Set, the output (*z*) of the network begins to ramp linearly toward a threshold (not to scale). The training target was a linear ramp (for other objectives see Supplementary Data Fig. 6) **(b)** The response profiles of randomly selected units in network (a) aligned to the time of Set. Many units exhibit temporal scaling. **(c)** Left: Network activity projected onto the first three principal components across all trials. Different traces correspond to trials with different durations (red for shortest to blue for longest). For each Cue input, the network engenders a pair of initial and terminal fixed points (circles; F_init_ and F_terminal_). Diamonds mark the state of the network along the trajectory 500 ms after Set. The Cue input moves the fixed points within an “Input” subspace. The corresponding trajectories for different intervals reside in a separate “Recurrent” subspace. Right: Rotation of the state space reveals the invariance of trajectories in the recurrent subspace. In the recurrent subspace trajectories traverse the same path at different speeds (see diamonds for different Cue inputs). **(d)** After training, the network accurately produced the intervals according to the presented Cue input. **(e)** A plot of the average speed in the recurrent neural network model as a function of the logarithm of the production interval (*Tp*). The speed was estimated from the rate of change of activity along the neural trajectory within the subspace spanned by the first three PCs. **(f)** Left: The spectrum of eigenvalues of the linearized dynamics near F_terminal_. Right: The spectrum of eigenvalues of a linear system 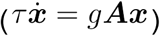. Applying a gain factor, *g*, to the transition matrix, *A*, causes eigenvalues to move radially (expand or contract) while the eigenvectors are preserved. The spectra corresponds to eigenvalues of an N-dimensional linear dynamical system with elements of *A* sampled from a distribution 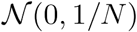. Decreasing the gain values from *g* = 1.0 (red) to 0.57 (blue) progressively decreases the magnitude of the eigenvalues and increases the effective time constants (*τ_eff_ = τ/g*). **(g)** Units in recurrent model were sorted based on their maximal activity when the network was near F_terminal_. The plot shows the maximum activity as function of Cue input. Vertical arrows mark two neurons, one with positive and another with negative activity, which are plotted in panel (h). **(h)** Stronger input drives units toward the saturation point of their nonlinear activation function where the shallowness of slopes leads to reduced gain of neural activity. This is true both for units with a positive response whose responses increased with Cue input (right), as well as units with a negative response, whose responses decreased with input drive (left). In all plots, different colors correspond to different intervals as shown by the color bar.

The network learned to generate the desired output function (Fig. 5d) and the activity of model neurons emulated the key features in MFC: response profiles of individual network units were heterogeneous, complex and temporally-scaled (Fig. 5b). Moreover, the speed of population dynamics directly determined the produced interval (Fig. 5c). The first set of networks we developed were trained to produce a linear output that was scaled according to the desired interval. To ensure that the scaling of network units was not merely a consequence of scaling in the output, we trained additional networks to produce temporally non-scaling output functions. However, even in the presence of non-scaling output functions, the recurrent model exhibited speed control in a scaling subspace and individual units continued to exhibit temporally scaled responses (Supplementary Fig. 6). The model was also robust with respect to how the input Cue encoded the desired interval. For example, the network exhibited the same scaling behavior when the Cue was changed to a brief pulse (Supplementary Fig. 6). The fact that the models captured subspace temporal scaling for various formulations of the input and for both scaling and non-scaling outputs demonstrated the generality and robustness of this solution in performing a flexible context-dependent timing task, and made the recurrent model suitable for uncovering potential underlying mechanisms.

We investigated how speed control emerges in recurrent neural populations (Fig. 5c) using a dynamical systems analysis ^65,66^ to study the evolution of activity in relation to neighboring fixed points in the state space. Importantly, we computed these fixed points in the presence of the input so that we could characterize the dynamics of the network in action. This analysis revealed that the network’s initial and terminal states were governed by a pair of input-dependent stable fixed points, F_init_ and F_terminal_. At the start of the trial, the Cue initialized the state of the network to an input-dependent fixed point, F_init_. Activation of the Set pulse drove the system away from F_init_ allowing the system to evolve toward F_terminal_ with a speed that was determined by the magnitude of the Cue input, which provided context (Fig. 5c,e).

This analysis revealed the complementary roles of the input drive and recurrent dynamics (Fig. 5c). The input drive set the position of the initial and terminal fixed points along a direction, which we refer to as the *input subspace*. Recurrent dynamics on the other hand, established a *recurrent subspace*, which determined the neural trajectory between the initial and terminal fixed points. These two subspaces emerged from different components of the network. The input subspace was governed by the direction specified by the input weights. In contrast, the recurrent subspace emerged from the constraints imposed by the recurrent weights. The two subspaces differed also in terms of their relationship to the scaling phenomenon. Within the input subspace, different intervals were associated with a change in the level of activity but did not exhibit scaling. This I change in level, in turn, controlled the speed by setting the position of the neural state along the axis of the input subspace. The recurrent space, on the other hand, did not control the speed but was responsible for the emergence of invariant trajectories and temporal scaling. The division of labour between these subspaces provides a remarkable and previously unsuspected explanation of why scaling and non-scaling signals might coexist within the same network. Non-scaling signals reflect the input that sets the speed, and scaling signals correspond to the evolution of activity with the desired speed. This organization predicts that MFC neurons with weak temporal scaling are likely recipients of relatively strong context-dependent input, and neurons with strong temporal scaling are more directly engaged in recurrent interactions. Finally, the model-based distinction between these two subspaces provides a theoretical basis for analyzing MFC responses within a scaling subspace, which corresponds to the recurrent subspace in the model.

We used the recurrent model to generate specific predictions about how activity in the scaling and non-scaling subspaces of MFC might correspond to behavior. Within the scaling subspace where neural trajectories were invariant, production intervals should be correlated with the speed of the dynamics. We demonstrated this earlier by analyzing production intervals in terms of speed within the space spanned by SCs (Fig. 3f). Within the non-scaling subspace where the activity putatively reflects the input, production times should be correlated with the average level – not speed – of neural activity. To test this novel model-based prediction, we asked whether production intervals could be predicted by the average MFC activity when projected onto the least-scaling subspace. We inferred the least-scaling direction from our scaling component analysis. SCs specified an orthonormal basis whose axes were ordered according to the level of scaling (Supplementary Fig. 7). Therefore, we used the last SC (SC9) as an estimate of the least-scaling direction, and compared production times to average level of MFC projected onto SC9. As predicted by the model, the average activity of the non-scaling components of MFC were indeed predictive of production times (Supplementary Fig. 8). This is a compelling result as it bears out a key prediction about an unsuspected relationship between cortical activity and behavior made by a model that was constrained only to produce behavior.

We emphasize that the recurrent model is fundamentally different from the oscillation and population-clock models since the former can support temporal scaling whereas the latter cannot. At a phenomenological level, the recurrent model superficially resembles the clock-accumulator model with a flexible clock since both exhibit temporal scaling. However, the two are conceptually and mechanistically different. The clock model can only generate ramping activity, and it does so by using the input (i.e., clock) to drive an integrator (e.g., a line attractor). The recurrent model does not perform any integration; instead, it uses an input to set the relaxation dynamics in the recurrent subspace toward a terminal fixed point. This difference allows the recurrent model to generalize the temporal scaling phenomenon across neurons with simple to highly complex response profiles.

## A potential neural mechanisms for speed control

To investigate the mechanisms that mediated the observed speed control, we analyzed the eigenvalues associated with the region in close proximity of the terminal fixed point, F_terminal_. In the vicinity of this fixed point, stronger inputs caused the eigenvalues to decrease systematically (Fig. 5f, left). In a linear dynamical system, such contraction in the eigenvalue spectrum corresponds to a systematic increase in the network’s effective time constants, *τ_eff_* (Fig. 5f, right). From this, we concluded that the action exerted by the input drive is equivalent to adjusting the system’s effective time constant in a flexible input-dependent manner. To gain insight into the mechanism that provides such powerful and modular control of time constants, we focused on a simplified recurrent network model composed of only two mutually inhibitory neurons that received a common input (Fig. 6a). The reasons for our choice of this model were twofold. First, this two-neuron network represents one of the simplest architectures whose behavior can be controlled by the interplay between common input and recurrent interactions (Fig. 6c). Second, previous work has demonstrated that adjustments of the common input in this model could alter its recurrent dynamics to either relax to a single fixed point with a specific time constant or act as an integrator with exceedingly long time constants ^67–69^. We reasoned that exploring the model’s behavior while between these two regimes might lead us to a mechanistic understanding of how effective time constant of a network can be flexibly adjusted.

**Figure 6.**
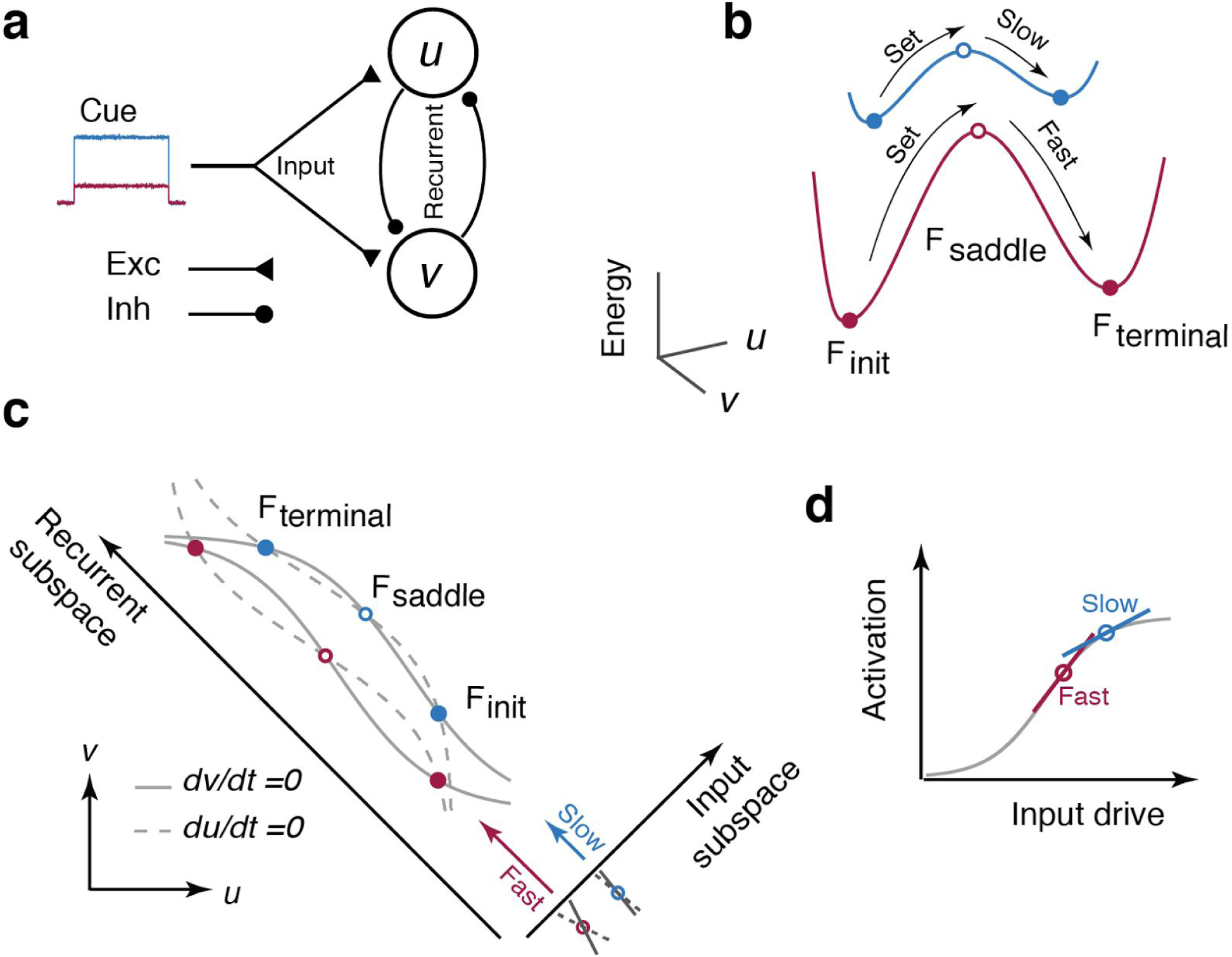
A simple two-neuron implementation of speed control. **(a)** Two inhibitory units (u and v) with recurrent inhibition receive a common excitatory input (Cue). **(b)** The energy landscape of the 2-neuron model. The network has a bistable energy landscape whose gradients depend on the strength of the Cue input. Stronger inputs (blue) lead to shallower energy gradients and vice versa (red). The Set pulse moves the state away from the initial fixed point (F_init_, filled circle) and over the saddle point (F_saddle_, open circle). The network then spontaneously moves toward the terminal fixed point (F_terminal_, filled circle). The speed of the movement toward F_terminal_ is relatively slow when the energy gradient is shallow (blue) due to stronger common input. **(c)** Phase plane analysis of the 2-neuron model. The two axes on the lower left correspond to the activity of the two neurons (u and v). The input is applied to both units and thus drives the system along the diagonal, which is labeled as “input subspace”. The input level moves the nullclines of the two units (du/dt = 0, dashed, and dv/dt = 0, solid) and adjusts the location of the three fixed points, (F_init_, F_terminal_ and the intermediate F_saddle_). The figure shows the two nullclines and the corresponding fixed points for two inputs levels (red and blue). Activation of Set moves the system along a “recurrent subspace” which is orthogonal to the input subspace. The proximity of nullclines (crosses below the Input subspace) controls the speed. When the input is stronger, the nullclines are closer, which causes the system to become slower. **(d)** Interaction of the input drive with the saturating nonlinearity of one unit. The action of the input upon the nonlinear activation functions moves the saddle point and controls the speed of the system. Stronger inputs push the neurons toward the shallower part of the nonlinear activation function, and moves the saddle point to slower regions of the phase plane causing the system to recurrent interactions to slow down.

When the two neurons receive balanced input (Cue), their interaction creates an energy landscape that engenders a pair of stable fixed points similar to the recurrent model that are separated by an energy barrier i.e., unstable fixed point (Fig. 6b). We analyzed the phase plane of the model (Fig. 6c) and verified that the input level can be used to create a continuum of *τ_eff_* as a function of the common input. This is analogous to the recurrent network model where activity along the input subspace served to control the speed. However, the two-neuron model helped us understand the underlying mechanisms in simple and intuitive terms: stronger input drives neurons toward their saturating nonlinearity where the slopes of activation functions are shallower (Fig. 6d). Regimes of shallow slopes reduce the neuron’s responsiveness to changes in input and lead to an increase in *τ_eff_*. In other words, the presence of single-neuron nonlinearities enable an input to exploit different slopes along the activation function. The slope would in turn determine the energy gradients (Fig. 6b) and set the speed with which responses change over time. What this simple network demonstrates is the importance of the action of the input on the neuron’s nonlinearities, which is the key factor in adjusting the speed.

Having established a low-level mechanism in the two-neuron model, we asked whether the same mechanism was operative in the recurrent network model, whose units are each equipped with a saturating nonlinearity analogous to the those found in the simple network. For the recurrent model, we analyzed the operating point of units as a function of the input drive near F_terminal_. Remarkably, for stronger inputs, units were systematically driven further toward their saturating nonlinearity (Fig. 5g,h), which is consistent with the mechanism of speed control in the simple network model. These results underscore a simple and powerful mechanism at the level of single neurons for controlling the speed of dynamics independent of the neural trajectory.

## Discussion

We found that motor timing in the MFC and caudate populations was governed by the rate of change of population activity as a function of time, which corresponds to the speed of dynamics. This property was evident at the single neuron level but was most pronounced in a scaling subspace of the population activity. In this subspace, the random fluctuations of speed predicted variability within each temporal context whereas systematic adjustments to the speed predicted behavioral flexibility across the two temporal contexts. These observations underline population speed within the scaling subspace as a key variable in motor timing behavior.

To gain insight into the principles of subspace speed control, we reverse-engineered a recurrent network model trained to perform a context-dependent timing task. The model demonstrated that flexible speed control is achieved through the interaction of two subspaces, an input subspace controlled by a context cue, and a recurrent subspace established by recurrent synaptic connections. These two subspaces served complementary functions that could be readily understood under the framework of dynamical systems. The input set the initial condition of the system and the recurrent subspace controlled the dynamics in the vicinity of that initial condition. In this framework, behavioral flexibility was conferred by the input’s ability to set the initial condition to different regions of the state space with different gradients (i.e., speeds).

To further understand how the context input is able to flexibly change the speed, we engineered a two-neuron model in which the input and recurrent subspaces could be represented as one-dimensional directions in the state space ^67–69^. This model highlighted the crucial role of single-neuron nonlinearities, revealing that adjustments of speed were governed by the interaction of input with these nonlinearities. This finding motivates several hypotheses regarding the structure and control of dynamics in the brain. For example, it suggests that many circuits and subcircuits might be able to adjust the speed of their dynamics independently and operate at different timescales ^70^. It also predicts that neuromodulatory effects and pharmacological treatments that interfere with the nonlinear response curve of individual neurons could alter the speed of cortical dynamics, as observations from numerous studies of interval timing might suggest ^71,72^.

Furthermore, our work has important implications for a wide range of behaviors in motor control, sensory anticipation and mental tracking where temporal flexibility is critical. The mechanisms we have identified provide a simple solution for how cortical circuits could encode behaviorally relevant variables in one subspace while using another subspace to adjust the speed of their dynamics. This would allow the same behavior to unfold along the same neural trajectory at different timescales. This is consistent with a study of the dynamics of sequential activation of striatal neurons in mice during a temporal bisection task ^73^, and studies of speed-accuracy tradeoff in decision making ^74^. Although these studies differ from ours substantially in terms of task demands and neural computations, the similarities in terms of neural dynamics raise the intriguing possibility that speed control might be a general principle in neural computations.

The source of the external input that adjusts the speed of cortical dynamics remains a pertinent and unresolved question. Several neural pathways could provide such input. One possibility is that the input arises from subpopulations of cortical neurons. This would be consistent with a recent study showing that neurons in parietal cortex encode the desired speed in their firing rates ^75^. Indeed, we found that MFC neurons that exhibited the least amount of scaling encode the speed in their average firing rates. Another possibility is that the input drive is provided by thalamic afferents, which is consistent with our recording from MFC-projecting thalamocortical neurons whose activity level varied systematically with produced intervals. Alternatively, a number of physiology and pharmacology studies have implicated dopamine in regulating timing behavior ^76,77^. Other neuromodulatory systems may also play a role in controlling cortical dynamics. Cortical dynamics are also known to depend on cellular properties such as those mediated by NMDA receptors, which are thought to facilitate the generation of stable slow cortical dynamics ^78^. The exact signalling pathways and underlying biophysical properties notwithstanding, the mechanisms that we have identified have the potential to explain how the brain flexibly controls cortical dynamics.

## Methods

Two adult rhesus monkeys (Macaca mulatta, a 6.5 kg female and 9.0 kg male, both 5 years old) were trained on a two-interval two-effector motor timing task. All surgical, behavioral and experimental procedures conformed to the guidelines of National Institutes of Health and were approved by the Committee of Animal Care at Massachusetts Institute of Technology.

## Behavior

The MWorks software package (<u>http://mworks-project.org</u>) running on a Mac Pro was used to deliver stimuli and to control behavioral contingencies. Visual stimuli were presented on a 23 inch monitor at a refresh rate of 60 Hz. Eye positions were tracked with an infrared camera (Eyelink 1000; SR Research Ltd, Ontario, Canada) and sampled at 1 kHz. A custom-made manual button, equipped with a trigger and a force sensor, was used to register button presses.

### Motor timing task

Each trial began with the appearance of two fixation cues (FCs), a circle at the center of the screen and a square 0.5 deg below the circle. The animal had to shift its gaze to the circle and the square informed the animal to hold its hand gently on the button. On each trial, one FC was colored and the other was white. The colored FC indicated the desired response effector (colored circle for saccade and colored square for button press). The color indicated the desired interval (red for 800 ms and blue for 1500 ms). We denote these four trial conditions by EL, ES, HL and HS where E and H refer to *Eye* and *Hand*, and S and L to *Short* (800 ms) and *Long* (1500 ms) intervals. After a delay period (500 - 1500 ms, uniform hazard), the saccade target was briefly presented 8 degrees to the left or right of the FC. For button press trials (colored square), the saccadic target was not relevant but was presented so that stimuli were consistent across trials. After another delay (500 - 1500 ms, uniform hazard), a 48 ms annulus (Set cue) flashed around the FCs cued the animal to start timing. Trials were aborted if the animal made premature eye or hand movements (before Set or long before the desired time). To receive reward, animals had to initiate a movement with the desired effector (cued by the colored FC) within a small window (“acceptance window”) around the desired interval (cued by the color of FC). The saccade responses had to land inside a circular window of radius 2.5 deg centered on the location of the extinguished target and had to be made directly (less than 33 ms after exiting the FC window). Button-press responses had to be made with the hand contralateral to the recorded hemifield ^79^. The production interval was measured from the endpoint of Set to the moment the saccade was initiated or the button was triggered. The width of the acceptance window was adjusted dynamically on a trial-by-trial basis and independently for the *Short* and *Long* conditions using a one-up one-down staircase procedure. As such, animals were rewarded for nearly half of trials (on average, 57% in monkey A and 51% in monkey D) for both temporal contexts. For trials that were rewarded, in addition to reward delivery, the color of the stimulus changed to green and an auditory clicking sound was simultaneously presented (Fig. 1a). Within the acceptance window, the magnitude of the reward scaled with accuracy.

## Electrophysiology

Animals were comfortably seated in a dark and quiet room. Each session began with an approximately 10-minute warmup period to allow animals to recalibrate their timing and exhibit stable behavior during electrophysiology recordings. Recordings were made through a craniotomy within a recording chamber while the animal’s head was immobilized. Structural MRI scans were used to aid in targeting regions of interest. Single- and multi-units responses were recorded using a 24-channel laminar probe with 100 μm or 200 μm interelectrode spacing (V-probe, Plexon Inc.). Eye position was sampled at 1kHz, and all behavioral and electrophysiological data were timestamped at 30 kHz and streamed to a data acquisition system (OpenEphys).

The dataset collected for this study included 1967 single- or multi-units recorded from the MFC, caudate and thalamus of two monkeys (Table 1), in which 69% (1351/1967) were tentatively single units. Neurons with firing rates less than 2 spikes per second during the timing epoch were excluded from subsequent analyses.

## Reversible inactivation

Injections were made with a microinjection pump (UMP3, World Precision Instruments) and a Hamilton syringe, which was connected to a custom 30G stainless steel injection cannula via a fused silica injection line (365μm OD, 100μm ID, Polymicro Technologies). In each injection session, we first established the animal’s baseline behavioral performance. Afterwards, we pressure-injected muscimol hydrobromide (5 μg/μL in saline) in the region of interest at a rate of 0.2 μL/min. In the MFC and caudate, a total of 2 μL was injected per session. In pilot inactivation experiments in the thalamus, we noticed that animals stopped performing the task after 2μL muscimol injection. To ensure animals would perform the task, the total volume of muscimol in the thalamus was reduced to 1.5 μL. The behavioral task was resumed 10 min after the the injection was completed. As a control, in separate sessions, sterile saline was injected following the same procedure. For each injection session, we compared performance between equal number of trials before and after the injection.

## Antidromic Stimulation

We used antidromic stimulation to localize thalamocortical MFC-projecting neurons. Antidromic spikes were recorded on a 24-channel electrode (V-probe, Plexon Inc.) in response to a single biphasic pulse of duration 0.2 ms (current < 500 uA) delivered to MFC via low impedance tungsten microelectrodes (100 - 500KΩ, Microprobes). The guide tube for the tungsten electrode was used as the return path for the stimulation current. Antidromic activation evoked spikes reliably at a latency ranging from 1.8 to 3 ms, with less than 0.2 ms jitter. The region of interest recorded in the thalamus was within 1 mm of antidromically identified neurons.

## Mathematical notation

Throughout the manuscript, we have used lowercase for scalars (*x*), bold and lowercase for vectors (***x***), bold and uppercase for matrices (***X***). Brackets were used for indexing values within vectors and matrices (***x***[*i*] and ***X***[*i*, *j*]) Subscripts were used for indexing a set of scalars (*x_i_*), vectors (***x****_i_*), or matrices (***X****_i_*). Subscripts were also used for specifying specific indices (***x***_*i*=1:*n*_). Superscripts were used for vectors to show projections onto a specific subspaces, for example, ***x***_*PC*(1:*k*)_, referring to a vector projected onto a the first *k* principal components. Curly brackets were used to indicate subset of conditions for which a variable is computed. For example, ***x***{*a* = *a*_0_; *b* = *b*_1_} refers to a vector computed for a subset of trials in which both *a* = *a*_0_ and *b* = *b*_1_ conditions were satisfied. The symbol ∪ is used to indicate data combined across a number of variables. For example 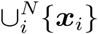 denotes data collected across a union of vectors ***x****_i_.* The symbol < ***x*** >*_i_* is used to show averaging of a vector ***x*** across *i.* Point functions were shown as lowercase (*f*(.)) regardless of whether they were applied to scalars or vectors.

## Data Analysis

All offline data processing and analyses were performed in MATLAB (MathWorks). Spiking data were bandpass filtered between 300 Hz to 7 kHz and spike waveforms were detected at a threshold that was typically set to 3 times the RMS noise. Single- and multi-units were sorted offline using a custom software, MKsort (https://sites.google.com/site/antimatt/software). The majority of the neurons were recorded in separate behavior sessions.

Estimating firing rates accurately is challenging when rates change dynamically and trials have different durations ^31,80^, which was the case in our data. Since our focus was on firing rates leading up to the movement, we aligned trials with respect to movement time (Fig. 2c). Additionally, for each condition, we discarded trials with *Tp*s that lay more than 3 standard deviations further from the mean (1.46% of trials). Firing rates were estimated by: 1) computing the peri-event time histogram (PETH) by averaging the spike counts per time bin, 2) using a 40 ms Gaussian kernel to compute smooth spiking density functions, and 3) z-scoring to minimize sampling bias due to baseline and amplitude differences across neurons.

To examine the relationship between firing rates and *Tp*s, we binned trials according to Tp and compared average firing rates for each bin. For the 800 ms interval, we used 7 bins centered on 740 to 860 ms every 20 ms, and for the 1500 ms, we used 9 bins centered on 1300 to 1620 ms every 40 ms. We denoted the overall average firing rate of a neuron as a function of time by *r*(*t*), average firing rate for a specific condition *c* (EL, ES, HL HS) by *r*(*t*; *c*), and average firing rate for a specific condition and a specific ***tp*** bin by ***r***(*t*; *c*, *tp_i_*). For population analyses, response vectors of individual neurons were organized into rows of a matrix denoted by ***r***(*t*; *c*, *tp_i_*).

To test if activity profiles could be described by a linear function (e.g. ramping activity), we compared 0 to 8th order polynomial fits to *r*(*t*) using cross-validation with randomized train and test sets. All neurons that were best explained by a polynomial of order 0 or 1 were considered linear so long as the fit explained at least 50% of variance. We also applied the same procedure allowing up to 200 ms offset from the beginning or end of the timing interval to ensure our results were robust.

## Compare the motor timing models at the level of single/multi-units

All model fitting was performed on the training set and the goodness of fit (R^2^) was quantified on the test set. In the clock-accumulator model with a flexible threshold, a linear ramp with fixed slope and different thresholds for different production intervals was fit to the response profile. In the clock-accumulator model with a flexible clock, the threshold was fixed and ramping rate was adjusted according to the interval. In the oscillation based models, sinusoidal functions or a sum of up to 4 different sinusoids were fit to activity profiles, in which the frequency, amplitude and phase for each sinusoid were free parameters. In the population clock, a single polynomial of up to 8th order was fit to the response profiles for both *Short* and *Long* contexts. For the temporal scaling model, the response profiles for the *Short* condition was used to find the best-fitting polynomial, and the temporally scaled version of the fitted functions were used to test the goodness of fit for *Long* trials.

## Scaling Subspace

We used a principal component analysis (PCA) as a first step to compute a low-dimensional and unbiased estimate of data. We found that the first 9 principal components (PCs) captured nearly 80% of the variance in the data (Fig. 3b, bottom). We therefore computed the scaling components (SCs) from data captured by the first 9 PCs, which was computed as follows:

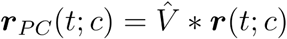 and 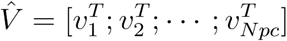 is the projection matrix, in which ***v_i_*** is the i^th^ PC direction. Therefore, the denoised activity across all conditions and time points ***r*** *_PC_*(*t*; *c*) is of size *N_PC_* × (*T* × *C*). we computed the corresponding scaled responses using our scaling procedure and denoted the result by 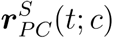. To find the scaling subspace, we solved an optimization problem that minimized the difference between average firing rates associated with different *Tp*s (e.g., *tp_i_* and *tp_j_*). We denote the corresponding projection by *U_SC_* and refer to its columns as scaling components (SCs). The resulting projection ***r****_SC_* can be computed as follows:

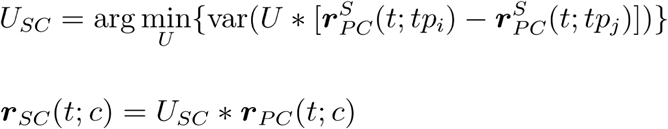

We hypothesized that the speed of activity in the scaling subspace predicts *Tp*. We computed the instantaneous speed in the scaling subspace from projections of responses on to the first three SCs as follows:

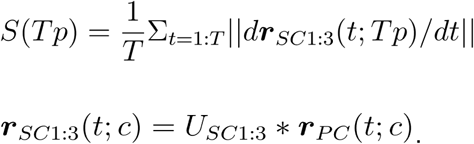

For each interval bin, we obtained an unbiased estimate of the relationship between speed and *tp* by resampling trials with replacement within each interval bin. The relationship between the average speed *S*(*Tp*) and production intervals was fitted in the log space by a linear function:

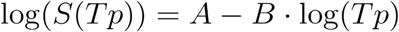

## Scaling index for population data

We quantified temporal scaling in single units, principal components (PCs) and scaling components (SCs) using a scaling index (*SI*) that represented a general measure of the degree of similarity between multiple response profiles associated with different intervals. *SI* was computed as follows: (1) trials were sorted based on production interval (*Tp*); (2) sorted trials were grouped into bins of similar *Tp*s (as described previously in Methods); (3) the first 9 PCs and the corresponding SCs for each bin were computed; (4) for each PC and SC, the index was computed as the coefficient of determination (*R*^2^) after the PCs and SCs were temporally scaled. This metric, which varies between 0 and 1, quantifies the degree to which each PC/SC undergoes temporal scaling for different *Tp*s.

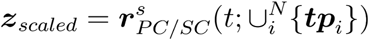

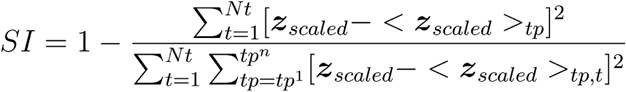

We evaluated the degree of scaling among populations in each region of interest by computing the scaling index for each PC and SC in those populations. Additionally, we computed the variance explained by each SC. Finally, to gain an unbiased estimate of the relationship between variance explained and scaling index, we computed these two metrics along randomly selected dimensions within the state space. This analysis revealed the full distributions of variance explained and scaling index and their relationship within the whole state space.

## Surrogate data for testing the scaling property

In order to assess the significance of scaling in firing rates recorded from the MFC, caudate and thalamus, we compared the *SI* computed for each region of interest to *SI* computed for data generated from a number of surrogate models that emulated various properties of the neural data with increasing levels of sophistication (Table 2). In what follows, we describe the four cumulative constraints that we considered for generating the surrogate data.

**Table 2.** Surrogate models

|  |  |
| --- | --- |
| Surrogate model 1 (S) | Smoothness constrained (S) |
| Surrogate model 2 (E+S) | Smoothness constrained (S)<br>Endpoints constrained (E) |
| Surrogate model 3 (D+E+S) | Smoothness constrained (S)<br>Endpoints constrained (E)<br>Dimensionality unconstrained (D) |
| Surrogate model 4 (C+D+E+S) | Smoothness constrained (S)<br>Endpoints constrained (E)<br>Dimensionality constrained (D)<br>Correlation constrained (C) |

### Surrogate model 1

We required the firing rates to have heterogeneous and nonlinear profiles while mimicking the same temporal smoothness as our data. To address this constraint, for each surrogate neuron, we sampled the response profiles (i.e., analogous to firing rates) from a zero-mean multinomial Gaussian Process (GP) prior ^81,82^. The covariance function between time points *t_i_* and *t_j_*, also known as the kernel function, *K*(*t_i_*, *t_j_*), was formulated as follows:

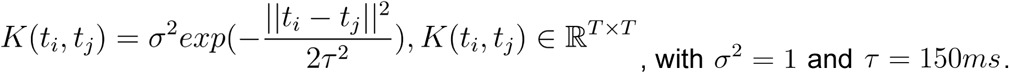

These parameters were chosen to mimic the average smoothness observed in response profiles of neurons. This constraint allowed us to examine whether the scaling indices associated with different regions of interest could be attributed to the the smoothness of firing rate functions. The surrogate model 1 was not worthy of consideration as i did not exhibit any scaling. But the smoothness constraint was used in the subsequent more constrained models.

### Surrogate model 2

The first model did not impose any structure among response profiles across the different produced intervals. In model 2, for each surrogate neuron, the starting point of the surrogate data was the same as the starting point of the real data and the endpoint of the Surrogate data was the same as the endpoint of the real data, but the starting point and end point of the real data were not necessarily the same. To satisfy this constraint, for each surrogate neuron, we first generated a response profile corresponding to the shortest produced interval, which we will refer to as the primary profile. We then constrained the response profiles associated with all longer produced intervals to match the primary profile at both the starting and terminal points. To do so, we sampled the remaining profiles from a conditional GP where the endpoint values were constrained to be the same as the primary profile. This constraint can be applied using GP regression analysis ^83^, where we derived the expectation and covariance of (*g*_2_,…,*g_tp_*_–1_) given the endpoints(*g*_1_, *g_tp_*). This model tested whether our scaling results could be attributed to a general level of similarity between smooth data with matched initial and terminal points.

### Surrogate model 3

The first two constraints ensured that the smoothness of the surrogate data and its starting/ending points corresponded to the recorded neuronal profiles. As a third constraint, we required the surrogate data to have the same dimensionality as the physiological data. To satisfy this constraint, we projected the surrogate data onto its principal components (PCs) and adjusted the variance according to the variance explained by PCs in neural data. This can be achieved by multiplying the data in the PC space by a diagonal matrix that scales each axis according to the desired variance. This model tests whether our scaling results could be attributed to data generated from a low-dimensional process with the same smoothness and matching initial and terminal points.

### Surrogate model 4

As a final constraint, we required the surrogate data generated for different produced intervals to exhibit the same coefficient of determination (R^2^) as the surrogate data generated from perfect scaling. To satisfy this constraint, for each surrogate neuron, we first sampled one instance from the GP for the shortest produced interval; i.e., the primary profile for that neuron. We then stretched the primary profile temporally to generate a set of perfectly scaled response profiles for all other produced intervals. We computed R^2^ as a measure of similarity between perfectly scaled responses. We then created samples from GP for other produced intervals that were matched in starting/ending points and had the same R^2^ as the perfectly scaled data. As a final step, we applied the same strategy as in surrogate model 3 to match the dimensionality of the surrogate and neural data. This model enabled us to test whether our scaling results could be attributed to the dimensionality of the data, given the same smoothness and similarity in response profiles achieved by matching temporal correlation as well as initial and terminal points.

We will refer to these models by the constraints they impose as follows:

The same analyses used to assess the neural responses were used to asses scaling in data generated from the surrogate models. This included (1) reducing the dimensionality to the first 9 PCs (which captured 80% of total variance of the physiological data), (2) computing the scaling index for each PC, (3) assessing the relationship between variance explained and scaling index for random projections of activity in the state space, (4) repeating the step for 200 times to obtain a distribution of scaling indices, and (5) comparing the distribution among different surrogate models and regions of interest.

## Recurrent Network Architecture

We constructed a firing rate recurrent neural network (RNN) model with *N* nonlinear units (*N =* 200). The network dynamics was governed by the following differential equation:

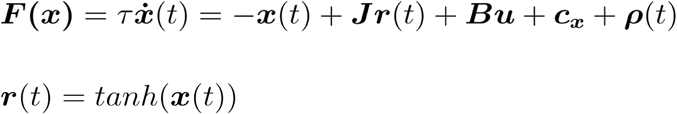

Variable ***x***(*t*) is an N-dimensional vector representing the activity of all the units. Variable ***r***(*t*) represents the firing rates of those units by transforming *x* through a *tanh* saturating nonlinearity. The time constant of each neuron was set to *τ* = 10*ms*. This value is different from *τ_eff_*, which emerges at the network-level. Variable ***c****_x_* is a vector representing a stationary offset the units receive, and ***ρ***(*t*) is a vector representing white noise *N*(0, .01) sampled at each timestep Δ*t* = 1*ms*. The recurrent connections in the network are specified by matrix ***J***, whose values, following previous work on balanced networks, are drawn from a normal distribution with zero mean and variance 1/N. The network receives a two-dimensional input *u* consisting of a context cue *u_c_*(*t*) and a transient Set pulse *u_s_*(*t*). The network received these inputs through synaptic weights ***B*** = [***b****_c_*, ***b****_s_*], which were initialized to random values drawn from a uniform distribution with range -1 to 1.

The context input, *u_c_*, represents the interval-dependent context cue input (color). The value of *u_c_* was was set to zero for 100 ms and then jumped to a graded value proportional to the length of one of 16 desired intervals distributed within a range 500 - 1700 ms. The offset of *u_c_* was sampled proportionally from the range 0.1 to 0.6 and was perturbed with Gaussian noise *N*(0, 0.025) at each Δ*t*. Increasing input noise did not qualitatively alter the network training solutions. The transient Set pulse *u_s_*(*t*) was active for 10 ms with magnitude 0.1 and zero elsewhere. On each training and test trial, the interval between the onset of *u_c_* and *u_s_*(*t*) was drawn from a uniform distribution with range (100 - 200 ms).

The network produced a one-dimensional output *z*(*t*), read-out by the summation of linear units with weights ***w****_o_* and a bias term *c_z_*. The output weights were initialized to zero at the start of training.

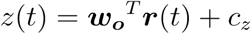

## Network Training

Multiple networks were trained with inputs that were presented randomly across trials as specified above. Networks were trained using backpropagation-through-time ^84^ and the Hessian-free (HF) method ^85,86^ was used to stabilize this. For HF, we used a preconditioner that utilized the diagonal of the generalized Gauss Newton matrix ^87^. We computed model parameters by minimizing a squared loss function between the network output ***z****_s_*(*t*) and a target function *f*_*s*_(*t*)^62^.

The objective least-squares function used during training was:

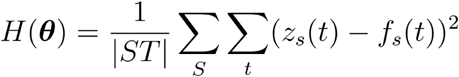

The target function *f_s_*(*t*) was computed for a subset of the trials and was only defined from the onset of the Set pulse till the length of the desired interval. For the duration of the Set pulse, the target function assumed a value of zero to prevent rapid transient switches in activity before the production stage.

Once training was complete, we generated a new set of randomized test trials, chose a threshold corresponding to the desired output (*z_s_ =* 1) and computed the times at which the network output crossed that threshold after Set. The time of threshold-crossing relative to Set was taken as the network’s produced interval (*Tp*). Network responses were also tested for intervals that interpolated within and extrapolated beyond the trained range. All networks exhibited good interpolation performance. Networks that exhibited good extrapolation performance within 150 ms beyond trained ranges were used for further analysis (90% of tested networks). In the main text, we report interpolated intervals that matched the monkeys’ response variability for both intervals.

We trained and tested multiple networks with different input profiles and different training objectives to check the consistency of the solution (Supplementary Data Fig. 6). In these networks, the scalar nature of the objective function was controlled by two parameters, *α* and *A*, to create a continuum of objective functions between a ramp (linear scaling) and a delta function (non-scaling). For example, with parameters A = 3 and alpha = 2.8, the above function approximates a scaled linear ramp of desired duration.

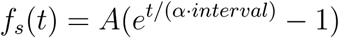

We tested additional networks with explicit non-scaling objectives and also with Cue inputs that did not remain constant throughout the trial. We have reported the results of two such variations in Supplementary Fig. 6, one in which the objective does not scale with interval duration (Supplementary Data Fig. 6a), and one in which the Cue input was a transient pulse (Supplementary Data Fig. 6d).

Following techniques adopted by Sussillo & Barak ^65^, we analyzed the rate of change of the network state to uncover its underlying dynamics. We computed ***q***(*x*) = 0.5 _*_ ∣***F***(***x***)∣^2^ to find the fixed and slow points of the system (corresponding to tolerances of *q* = 1*e* – 18 and 1*e* – 5 respectively). We analyzed the local dynamics around fixed points by performing an eigendecomposition of the corresponding linearized system. Fixed points associated with no positive eigenvalues were classified as stable fixed points. The unstable fixed points (*F_saddle_*) that we detected in some networks were usually associated with one unstable mode and often helped mediate the transition of trajectories after Set towards F_terminal_.

We projected activities of neurons on each trial onto the first three principal components of the total activity covariance matrix (across all trials) to obtain state space trajectories (Fig. 5c). Fixed points were projected onto the same basis for visualization purposes. For various networks, the mean squared speed of network trajectories during interval production over the first three PCs was used to relate dynamics to *Tp* (Fig. 5, Supplementary Figure 6). We additionally computed the scaling components (SCs) for the activity of various networks (Supplementary Fig. 7) The speed of the trajectory within the scaling subspace was calculated based on projection onto the first 3 SCs (Supplementary Data Fig. 7).

## Two inhibitory-neuron model

Following previous work ^69^, we engineered a two-neuron model that captured the key intuition behind speed control in the interval timing task. The network consists of two mutually inhibitory neurons (*u*, and *v*), which receive a common input representing the contextual cue, denoted as *θ* (Fig. 6a). The time-varying firing rates of the model neurons are governed by a pair of nonlinear differential equations as follows:

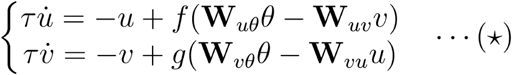

Model structure and parameters were as follows:

|  |  |  |
| --- | --- | --- |
| $\mathbf{W}_{ij}$ | $\mathbf{W}_{u\theta} = \mathbf{W}_{v\theta} = 6$<br>$\mathbf{W}_{uv} = \mathbf{W}_{vu} = 6$ | Magnitude of synaptic weight from input j to unit i |
| $\theta$ | $\theta = 0.6, 0.8$ | Balanced input to $u$ and $v$ |
| $\tau$ | 100 ms | Time constant of each unit,<br>$\Delta t = 10ms$ (time step) |
| $f(x), g(x)$ | $\frac{1}{1 + \exp(-x)}$ | Activation function for $u$ and $v$ respectively |

The nullcline for each variable is defined as the set of all the states where the derivative of the variable vanishes. For example, solving for 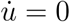 provide a solution of the kind *u* = *f* (*θ*, *v*), which represents the nullcline for *u* (Fig. 6c). The intersection of the nullclines pertaining to each variable represent a point where variables do not change, i.e., fixed points (FPs). For a range of synaptic weights (**W***_ij_*) and input levels (*θ*), the system has two stable FPs that are separated by an unstable one. Increasing *θ* systematically shifts the position of the two nullclines (Fig. 6c). It also_reduces the energy gradient between the FPs (Fig. 6b) causing the system to slow down.

We numerically solved the coupled equations (⋆) to compute the system’s response dynamics. The energy (E) of the system was defined as the path integral over a vector field: 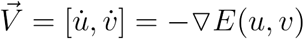. This was computed numerically by integrating the speed along a deterministic trajectory from the unstable FP to the stable ones using the equations above (⋆) to numerically simulate the relaxation of the activity towards the final FP.

## Acknowledgements

We thank Michale Fee, James DiCarlo, and Robert Desimone for comments on the manuscript, and David Sussillo for advice on modeling. M.J. is supported by NIH (NINDS-NS078127), the Sloan Foundation, the Klingenstein Foundation, the Simons Foundation, the Center for Sensorimotor Neural Engineering, and the McGovern Institute. D.N. is supported by the Rubicon Grant (2015/446-14-008) from the Netherlands Scientific Organization (NWO).

---

**Table 1.**
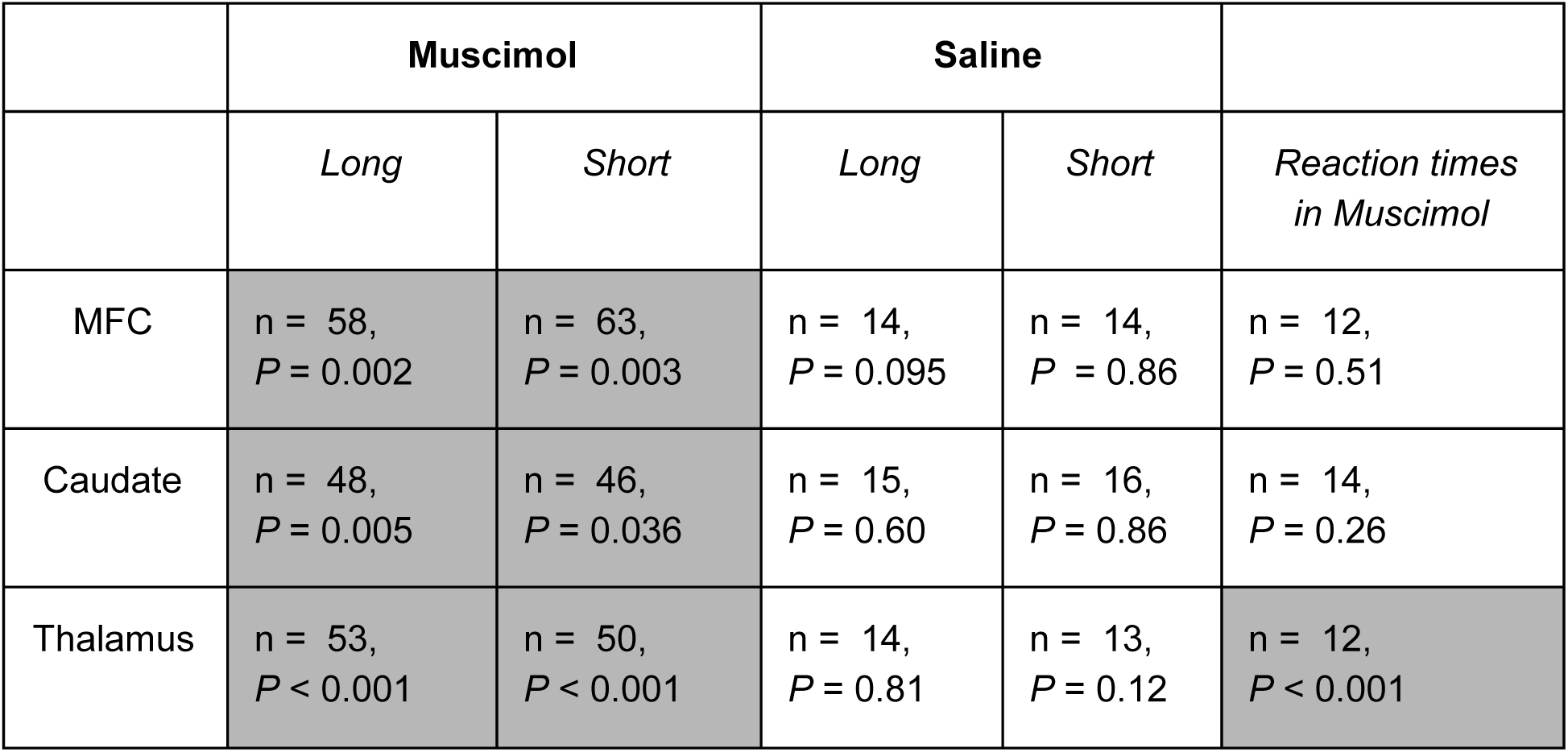
Effects of muscimol inactivation in the three brain areas. In assessing the significance of inactivation in the motor timing experiment (second and third columns from left), we used one-tailed paired-sample Student’s *t*-tests to examine whether the muscimol caused an increase in RMSE (HO: no increase in RMSE). The same test was used for the saline injection experiments (third and fourth columns from left). Statistical tests were done based on distributions of average RMSEs derived from trials before and after muscimol injection. Each pair of average RMSE values were computed from a pair of 50 random trials from before and after injection. The sampling was made without repeats to ensure trials were not counted twice. In assessing the effect of muscimol on reaction times (RTs) in the memory saccade task (the rightmost column), we used a two-tailed Student’s *t*-tests to examine whether the muscimol caused a change in reaction time (HO: no change in RT). Statistically significant effect are marked in gray. There was a change in RT after muscimol inactivation in the thalamus. This suggests that the region influenced by the inactivation of thalamus could also have a role in the memory saccade task.

**Supplementary Figure 1.**
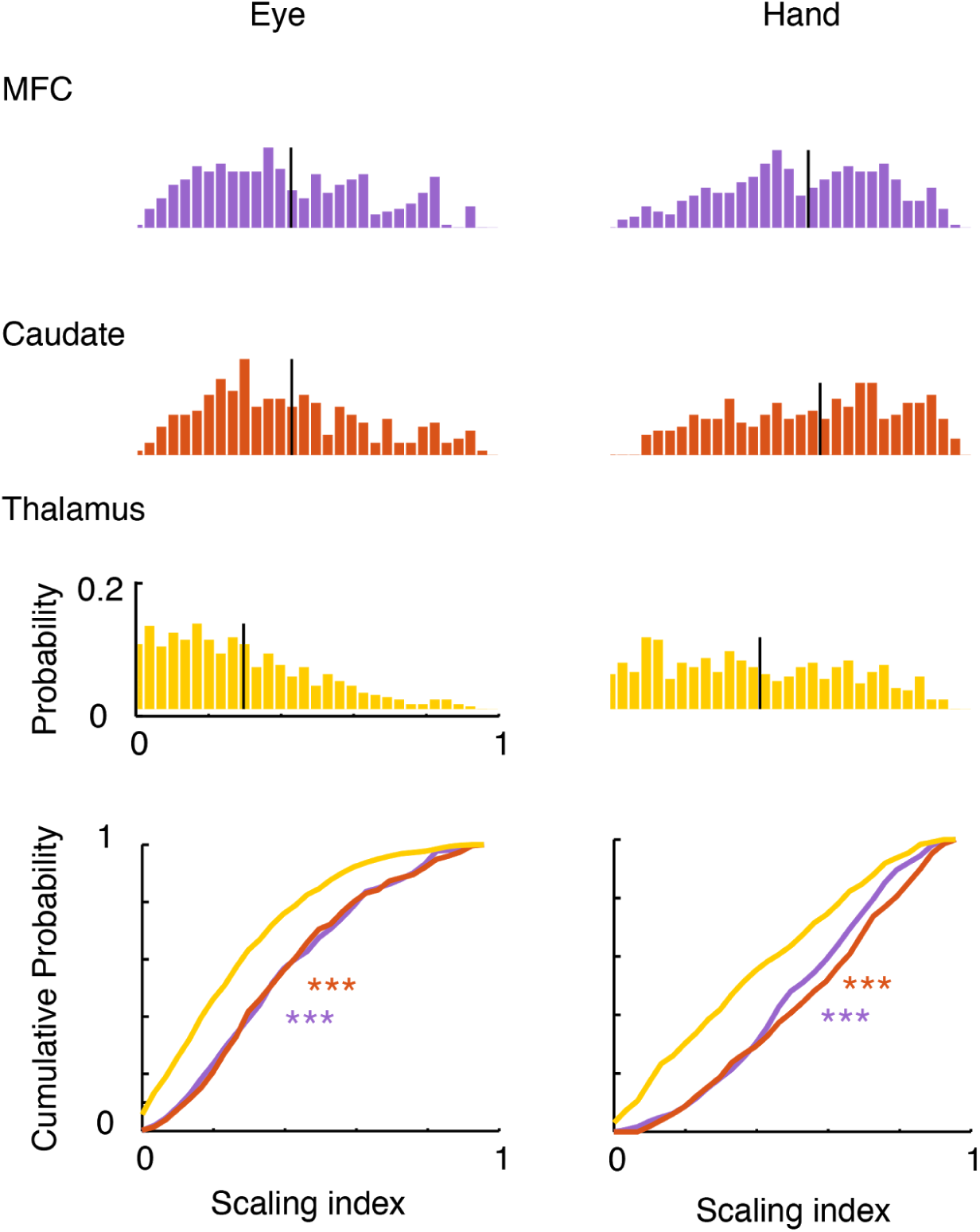
Scaling index of single neurons in different brain areas. Histograms on the top show the normalized distribution of scaling index for individual neurons in the MFC (n = 416), caudate (n = 278), and thalamus (n = 846) for the *Eye* (Left) and *Hand* conditions (Right). The bottom panel shows a comparison of the cumulative probability distribution of scaling index across the three areas. The thalamus shows a predominance of smaller scaling index value (one-tailed two sample *t*-test, *P* < .001).

**Supplementary Figure 2.**
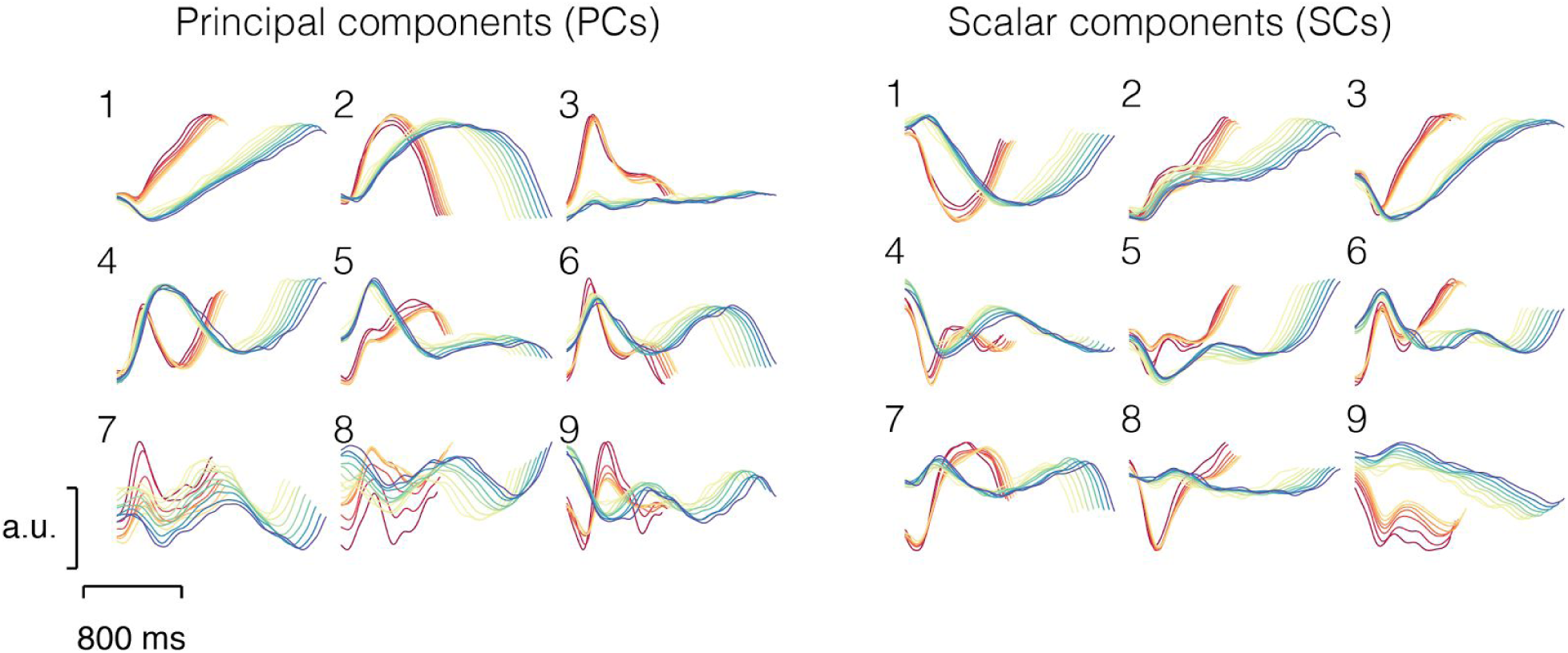
The time course of PCs (left) and SCs (right) for MFC data. First 9 PCs over the course of the production interval (abscissa) that explain 80% of variance in the MFC data in decreasing order of variance explained (left) for Short (warm colors) and Long (cool colors) intervals. First 9 SCs, obtained for the same data, in decreasing order of scaling (right, see Methods).

**Supplementary Figure 3.**
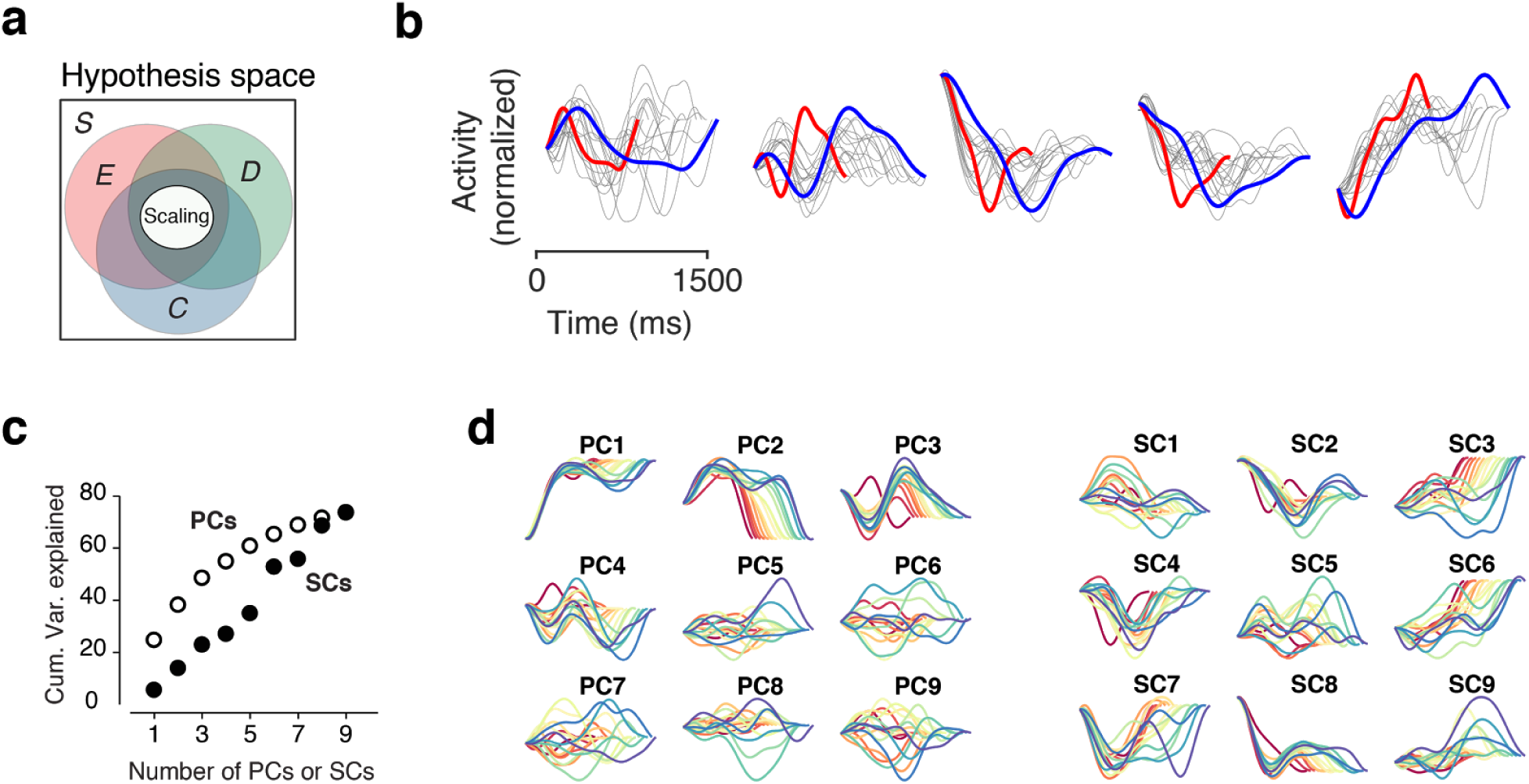
Analysis of scaling with surrogate data. **(a)** Venn diagram showing the various constraints considered for non-scaling models. All surrogate data was generated from a Gaussian Process (GP) with the same level of temporal smoothness (white rectangle, *S*) as the data. We considered three additional constraints to make the surrogate data more similar to neural data without an explicit requirement for scaling. One constraint required responses for all production intervals to be at the same level at the time of Set and at the time of response. We refer to this constraint as endpoint matching (red circle, *E*). Another constraint required that the dimensionality of the surrogate data match the neural data, and additionally the variance explained by each principal component (PC) be matched. We refer to this constraint as dimensionality matching (green circle, *D*). Finally, we considered a constraint that required the collection of responses for different production intervals to have the same correlation (quantified as R^2^) as expected from perfect scaling. We refer to this constraint as correlation matching (blue circle, *C*). We created surrogate data for each constraint and for various combination of constraints, and compared the scaling properties to the original data. Note that each constraint characterized a superset of the scaling hypothesis. **(b)** Example traces showing the procedure for generating the surrogate data in the *C+D+E+S* model for 5 randomly selected surrogate units aligned to the time of Set. We first sampled a *Short* trace (red) from a Gaussian process. The trace in blue corresponds to the perfectly scaled version of the red trace and is not a sample from the surrogate model. The surrogate data were generated using a constrained Gaussian Process (GP) prior as follows: the response for the shortest production intervals (red) was sampled from a GP with the same level of temporal smoothness as the neural data. The corresponding response with perfect scaling was generated by linear scaling (shown in blue). Note that the trace in blue is not a sample from the GP and is therefore, not part of the surrogate data. The gray traces correspond to the surrogate data. To generate the surrogate data, we drew samples from the Gaussian process that satisfied several criteria. First, the starting point as well as the ending point of every gray trace had to be perfectly matched to the starting point and ending point of the perfectly scaled blue trace. Second, across the population of surrogate data, the dimensionality had to match observed neural data. Finally, the correlation between every gray trace and the red trace was the same as the correlation between the red and blue trace. In this way, every sample of GP (gray traces) matched the smoothness, endpoints, dimensionality and correlation as the real data (i.e., *C+D+E+S* model). **(c)** Cumulative percentage variance explained by PCs and SCs for the surrogate data generated from the non-scaling *C+D+E+S* model. **(d)** The first 9 principal components of population activity (PC, left) and the corresponding 9 scaling components (right, SCs) plotted as the function of time from Set for the non-scaling *C+D+E+S* model. Note that PCs and SCs are based on the surrogate data (gray traces in panel b) – not the perfectly scaled data (blue traces in panel b).

**Supplementary Figure 4.**
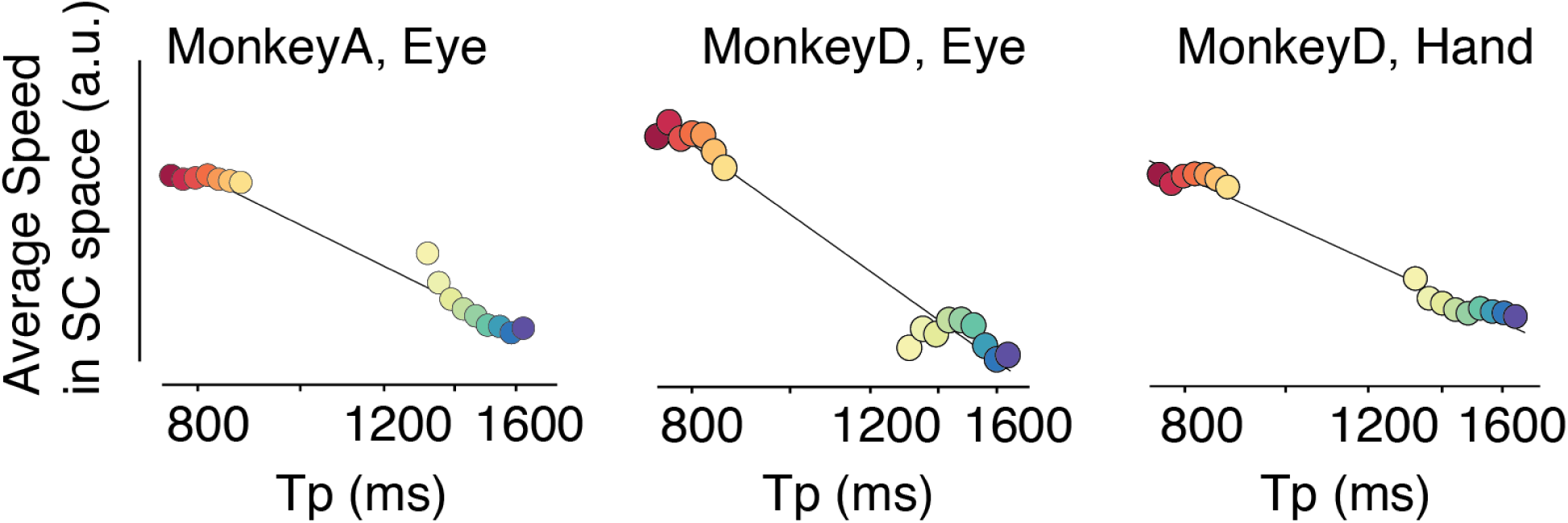
Relationship between MFC population activity and behavior. The speed of neural trajectory in MFC within the scaling subspace spanned by the first 3 SCs predicted *Tp* across both *Short* and *Long* conditions on a trial-by-trial basis. The case of hand trials for Monkey A was shown in Fig. 5d. Here, all the other conditions are shown.

**Supplementary Figure 5.**
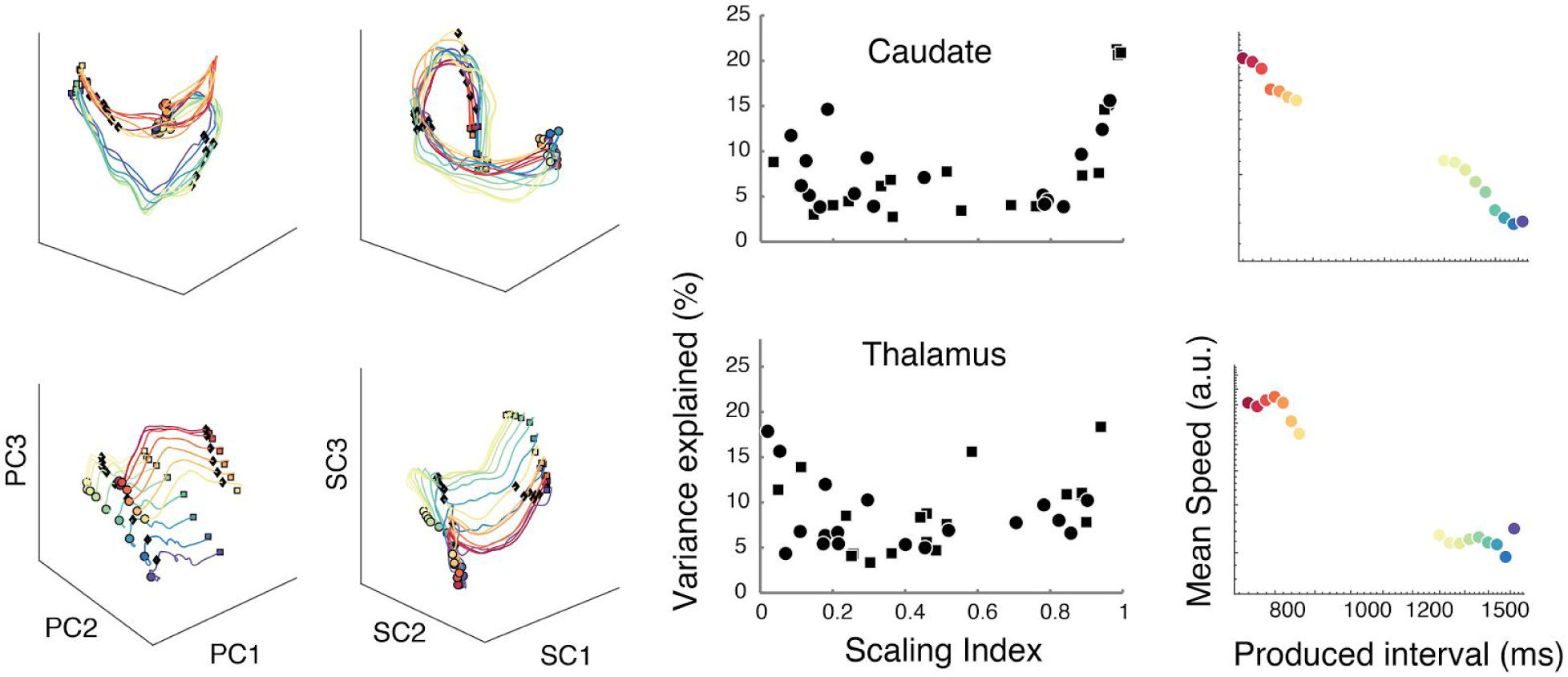
Analysis of scaling at the population level in the caudate (top) and thalamus (bottom). Left column: Population activity profiles projected onto the first 3 principal components (PCs). Activity profiles associated with different produced intervals for *Short* and *Long* conditions are plotted in different colors (same color scheme used throughout the paper). The state at 700 ms after Set is shown along the trajectories (diamond). Second column from the left: Population activity projected onto the first 3 scaling components (SCs). Activity spanned by the first 3 SCs overlap for different intervals in the caudate but not in the thalamus. Third column from left: Variance explained for individual SCs as a function of scaling index. Right column: The speed of neural trajectory within the scaling subspace spanned by the first 3 SCs.

**Supplementary Figure 6.**
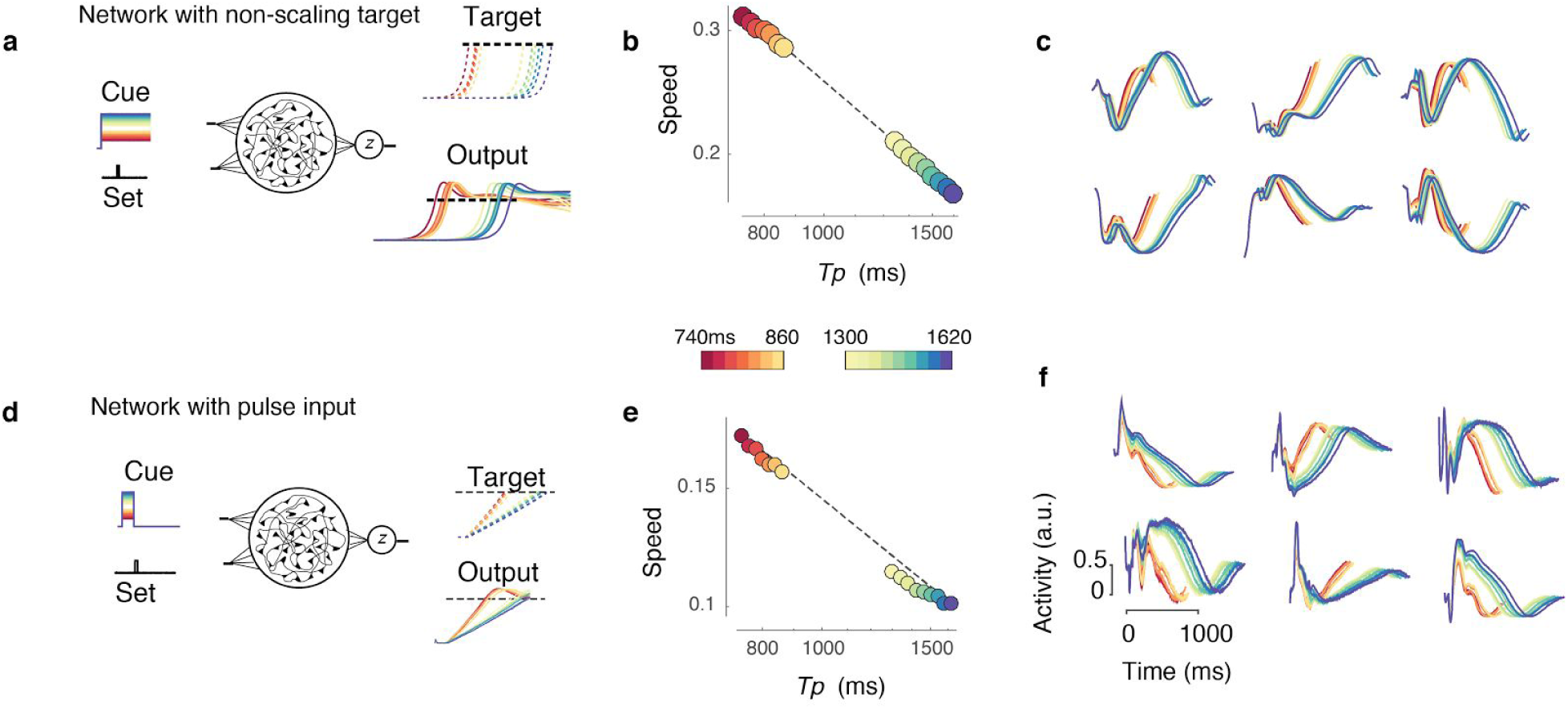
Alternative recurrent neural network models. **(a)** A recurrent neural network (RNN) trained to use an interval-dependent Cue input to produce time intervals flexibly. The network was trained using a non-scaling exponential objective with a fixed time constant. **(b)** The speed of dynamics measured within the space spanned by the first three PCs predicted *Tp* across both *Short* and *Long* conditions. **(c)** The response profiles of randomly selected units in the network (a) aligned to the time of Set. **(d)** A RNN trained to use a brief pulse as the interval-dependent Cue input to produce time intervals flexibly. The network was trained using a linear ramping objective like the network in the main text. **(e)** The speed of dynamics predicted *Tp* across both *Short* and *Long* conditions. **(f)** The response profiles of randomly selected units in network (d) aligned to the time of Set.

**Supplementary Figure 7.**
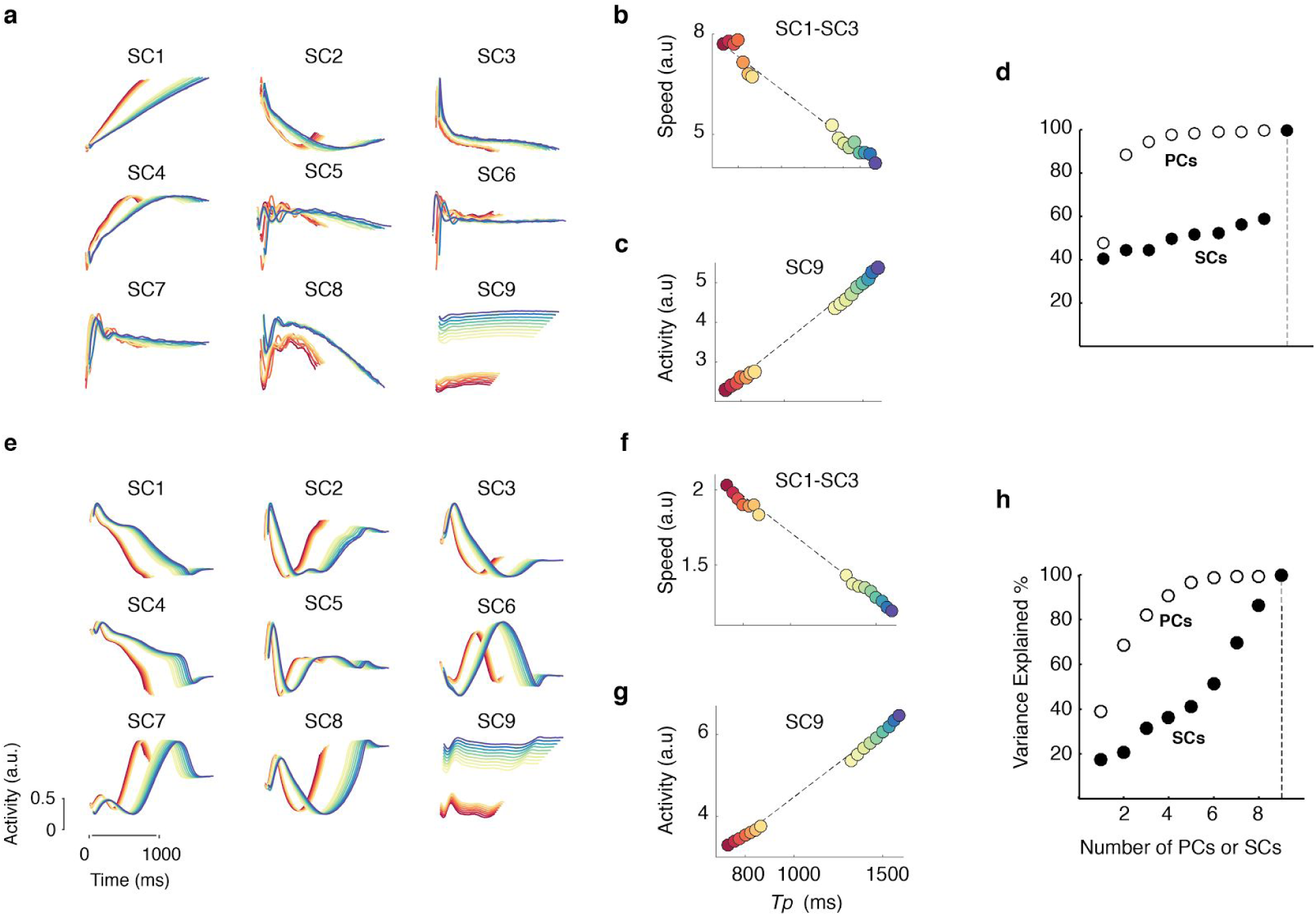
Temporal scaling in the recurrent network model. **(a)** The first 9 scaling components (SCs) of the population activity in the RNN with a scaling output function (Fig. 5). The early SCs correspond to the recurrent subspace, and the last SC represents the input subspace. **(b)** The average speed of population activity in the subspace spanned by the first 3 SCs is predictive of both within context and across context variations in *Tp.* **(c)** The average firing rate of the population activity projected onto the last SC is also predictive of *Tp.* **(d)** Cumulative percentage variance explained by PCs (white) and SCs (black). The dashed vertical line correspond to the 9th component. **(e-h)** Same as a-d, for a network that was trained for a non-scaling exponential output objective function (Supplementary Fig. 6d).

**Supplementary Figure 8.**
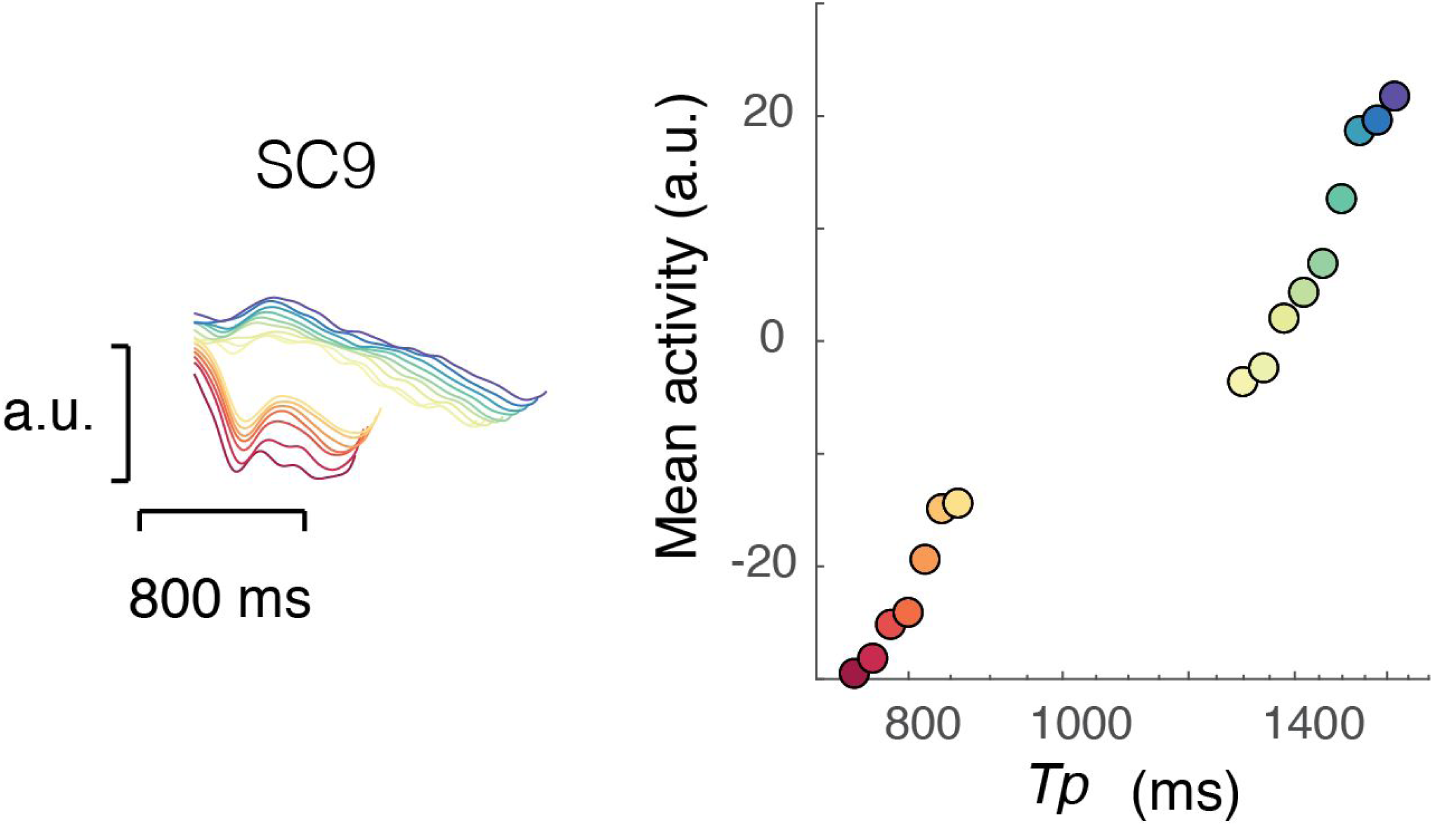
Non-scaling population activity in MFC. Left: The time course of the SC9 (the least scaling component) across conditions. Right: The average firing rate of population activity projected onto SC9 (left), also the putative input subspace, increases with produced intervals (*Tp*). This is consistent with the hypothesis that the average firing rate in the non-scaling subspace controls speed. Based on the recurrent network model, this subspace likely reflects the input drive to MFC.

